# Genetic structure correlates with ethnolinguistic diversity in eastern and southern Africa

**DOI:** 10.1101/2021.05.19.444732

**Authors:** Elizabeth G. Atkinson, Shareefa Dalvie, Yakov Pichkar, Allan Kalungi, Lerato Majara, Anne Stevenson, Tamrat Abebe, Dickens Akena, Melkam Alemayehu, Fred K. Ashaba, Lukoye Atwoli, Mark Baker, Lori B. Chibnik, Nicole Creanza, Mark J. Daly, Abebaw Fekadu, Bizu Gelaye, Stella Gichuru, Wilfred E. Injera, Roxanne James, Symon M. Kariuki, Gabriel Kigen, Nastassja Koen, Karestan C. Koenen, Zan Koenig, Edith Kwobah, Joseph Kyebuzibwa, Henry Musinguzi, Rehema M. Mwema, Benjamin M. Neale, Carter P. Newman, Charles R.J.C. Newton, Linnet Ongeri, Sohini Ramachandran, Raj Ramesar, Welelta Shiferaw, Dan J. Stein, Rocky E. Stroud, Solomon Teferra, Mary T. Yohannes, Zukiswa Zingela, Alicia R. Martin, NeuroGAP-Psychosis Study Team

## Abstract

African populations are the most diverse in the world yet are sorely underrepresented in medical genetics research. Here, we examine the structure of African populations using genetic and comprehensive multigenerational ethnolinguistic data from the Neuropsychiatric Genetics of African Populations-Psychosis study (NeuroGAP-Psychosis) consisting of 900 individuals from Ethiopia, Kenya, South Africa, and Uganda. We find that self-reported language classifications meaningfully tag underlying genetic variation that would be missed with consideration of geography alone, highlighting the importance of culture in shaping genetic diversity. Leveraging our uniquely rich multi-generational ethnolinguistic metadata, we track language transmission through the pedigree, observing the disappearance of several languages in our cohort as well as notable shifts in frequency over three generations. We find suggestive evidence for the rate of language transmission in matrilineal groups having been higher than that for patrilineal ones. We highlight both the diversity of variation within the African continent, as well as how within-Africa variation can be informative for broader variant interpretation; many variants appearing rare elsewhere are common in parts of Africa. The work presented here improves the understanding of the spectrum of genetic variation in African populations and highlights the enormous and complex genetic and ethnolinguistic diversity within Africa.

## Introduction

Humans are believed to have originated in Africa, resulting in more genetic variation on the African continent than anywhere else in the world; the average African genome has nearly a million more genetic variants than the average non-African genome(Consortium and The 1000 Genomes Project Consortium, 2012). Africa is also immensely culturally and ethno-linguistically diverse; while the rest of the world averages 3.2 to 4.7 ethnic groups per country, African countries have an average of greater than 8 each and account in total for 43% of the world’s ethnic groups(Fearon, 2003). Despite this diversity, African ancestry individuals are sorely underrepresented in genomic studies, making up only about 2% of GWAS participants(Popejoy and Fullerton, 2016; Sirugo et al., 2019). Furthermore, the vast majority of African ancestry populations currently represented in genetic studies are African Americans or Afro-Caribbeans (72-93% in the GWAS catalog and ≥ 90% in gnomAD) with primarily West African ancestral origins(Martin et al., 2018). These resources thus currently leave out the substantial diversity from regions of Africa that would be informative for human genetics.

Populations underrepresented in genetic studies contribute disproportionately to our understanding of biomedical phenotypes relative to European ancestry populations. Despite their paltry representation in GWAS, African ancestry populations contribute 7% of genome-wide significant associations(Martin et al., 2018; Morales et al., 2018). African population genetic studies are especially informative given their unique evolutionary history, high level of genetic variation, and rapid linkage disequilibrium decay(Tishkoff and Verrelli, 2003). This Eurocentric bias in current genomics studies and resources also makes African descent individuals less likely to benefit from key genomic findings that do not translate fully across populations, contributing to health disparities(Martin et al., 2019). In this study, we better characterize the immense genetic and ethnolinguistic diversity in four countries in eastern and southern Africa, offering insights into their population history and structure. Data are from 900 genotype samples that are part of the Neuropsychiatric Genetics of African Populations-Psychosis study (NeuroGAP-Psychosis), a major research and capacity building initiative in Ethiopia, Kenya, South Africa, and Uganda(van der Merwe et al., 2018; Stevenson et al., 2019)

Genetic variation in Africa has been previously described as following not only isolation-by-distance expectations, but as being influenced by multiple interconnected ecological, historical, environmental, cultural, and linguistic factors (Baker et al. 2017; Uren et al. 2016; Henn et al. 2012; Henn et al. 2016; Sikora et al. 2011; Creanza et al. 2015; Kolodny et al. 2016; Creanza and Feldman 2016). These factors capture distinct variation from that tagged by genetics and can be informative for understanding population substructure. Better characterization of the ethnolinguistic composition of these samples is a key initial step towards running well-calibrated statistical genomics analyses including association studies. If ethnolinguistic variation tags additional structure than that captured by geography, explicit incorporation of relevant cultural information into such analyses tests may be the optimal strategy. In addition to the covariation of culture and genetics(de Filippo et al. 2011; Karafet et al. 2016; Barbieri et al. 2013), individuals’ cultural environments influence how phenotypes are expressed and whether assortative pairing impacts the distribution of traits(Coelho et al. 2009; Uchiyama et al. 2021; Creanza et al. 2017). We measure how ethnolinguistic culture has changed in parallel to and independently of genetics, which provides a foundation for the study of phenotypes of medical interest. In this study, we explore the genetics of the NeuroGAP-Psychosis dataset, which comprises five collection sites across four countries in Africa, and how individuals’ cultural affiliations and languages are related to genetic variation across the continent. We also explore ongoing linguistic changes and consider the impact they will have on the genetics of Africa.

## Results

### Genetic Population Structure and Admixture

We compared the ancestral composition of our samples relative to global reference data from the 1000 Genomes Project, Human Genome Diversity Panel, and the African Genome Variation Project (AGVP) to see the full breadth of genetic diversity(Bergström et al. 2020; Auton and Salcedo 2015; Gurdasani et al. 2015). Most NeuroGAP-Psychosis samples appear genetically similar to their geographically closest reference samples when compared to global datasets (**Figure 1, Supplementary Figures 1**). However, large amounts of admixture is visible within some individuals, particularly among South African individuals (*Supplemental Information*). In South Africa, some individuals cluster wholly within the European reference cluster; this is expected based on the demographic composition of Cape Town, where these samples were collected, which is home to a substantial fraction of people of Dutch ancestry (Afrikaners) and individuals of mixed ancestry(Bergström et al., 2020; Chimusa et al., 2015; Pickrell et al., 2014; Sikora et al., 2011; Uren et al., 2016).

**Figure 1.**
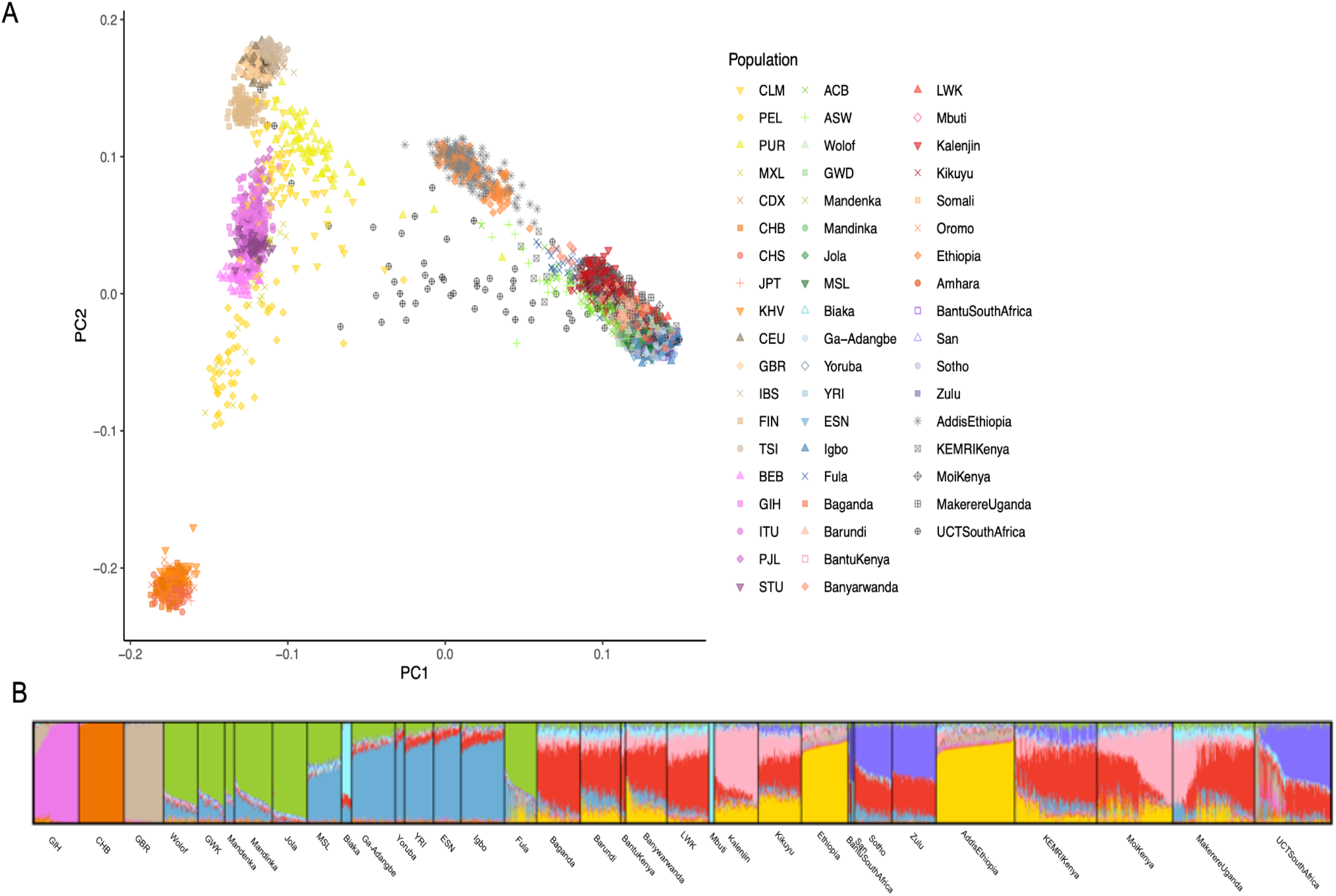
Genetic and admixture composition of the NeuroGAP-Psychosis samples against a global reference. A) First 2 principal components showing NeuroGAP-Psychosis samples as projected onto global variation of the full 1000 Genomes, HGDP, and AGVP. While most samples fall on a cline of African genetic variation, some South African samples exhibit high amounts of admixture and European genetic ancestry. Color scheme for global PCA plot: Latinx American - yellow; East Asian - dark orange; European - tan; South Asian - fuschia; West African - green/blue; East African - red/orange; South African - purple; NeuroGAP-Psychosis collections - gray. B) ADMIXTURE plot at best fit k (k=10) of all African samples as well as three representative non-African populations from the 1000 Genomes Project. The GIH, CHB, and GBR were included to capture South Asian, East Asian, and European admixture, respectively. Individuals are represented as bar charts sorted by population, and ancestry components for each person are visualized with different colors. A key describing the country of origin for all populations can be found in Supplemental Table 1.

We additionally investigated the degree of admixture within samples and how genetic groups cluster in the data. We ran ADMIXTURE(Alexander et al. 2009), which partition genetic variation into a given number of distinct genetic clusters. This helps to visualize the groups that are most genetically distinct from one another, as each additional component can be thought of as representing the next most differentiated ancestry component in the data, akin to principal components analysis (PCA). We identified the best fit *k* value, using five-fold cross validation, to be 10 using a tailored global reference.

Examining the ancestry composition at the best fit *k*, we identify several ancestry components unique to areas within continental Africa (**Figure 1B**, Supplementary Figure 1). Notably, several such components, including those unique to Ethiopia (yellow), West Africa (blue), and South Africa (purple) appear at earlier values of *k* than that separating South Asians from East Asians and Europeans (fuschia from tan and orange). This suggests a high level of genetic differentiation between areas of the African continent rivaling that between those out-of-Africa continental ancestries, as has been previously demonstrated. We also note that Ethiopian participants have evidence of Eurasian admixture, possibly related to historical back-migration into the African continent(López et al., 2021a; Pagani et al., 2012, 2015; Pickrell et al., 2014).

Projecting our samples onto PC space generated from only African reference samples, the top two principal components (PCs) separate geography, and more specifically east-west and north-south patterns of variation within Africa (**Figure 2**), mirroring isolation by geographic distance in human genetic data. At higher PCs, however, there is fine-scale structure in the data separating different geographically proximal groups within the East African individuals, shown in red. We thus focus our deeper examinations into the East African samples to assess potential drivers of this differentiation (**Supplementary Figures 2-6**). Clear structure is visible in the data to PC8, with higher PCs resolving substructure within geographic regions. For example, two clusters are evident among PCs within participants enrolled in the study from Moi University in Kenya, who tend to speak distinct languages in the Afro-Asiatic and Niger-Congo families (**Supplementary Figure 4**). For a detailed discussion of genetic variation within each country see the *Supplementary Information*.

**Figure 2.**
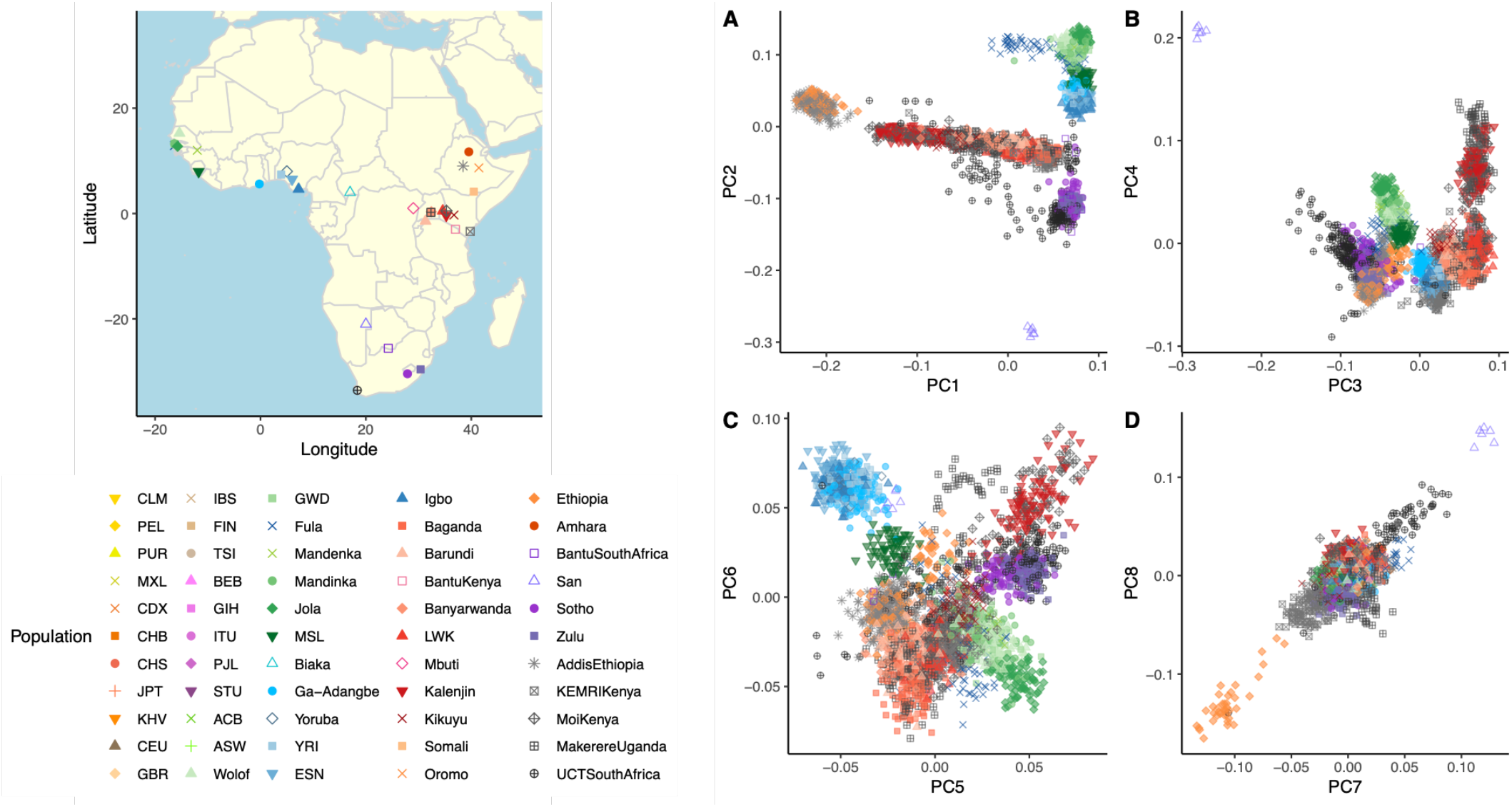
Genetic composition of subcontinental African structure in the NeuroGAP-Psychosis samples. A-D: PCA plots for PCs 1-8 with an African reference panel. A map of collection locations is shown to the left of PCA plots. Points are colored by region to assist in interpretation: green - west, blue - west central/central, red - east (orange - Ethiopia), purple - south. See Supplementary Figures 2-6 for plots highlighting each cohort individually.

### Self-reported Population Composition

Across samples with self-reported ethnolinguistic information, we observe 62 primary ethnicities and 107 primary languages in the 960 NeuroGAP-Psychosis samples. We also find that languages have shifted in frequency over time, with English increasing in reporting frequency in the current generation, and several grandparental languages disappearing in our dataset (**Figure 3; Supplementary Figure 7**.

**Figure 3.**
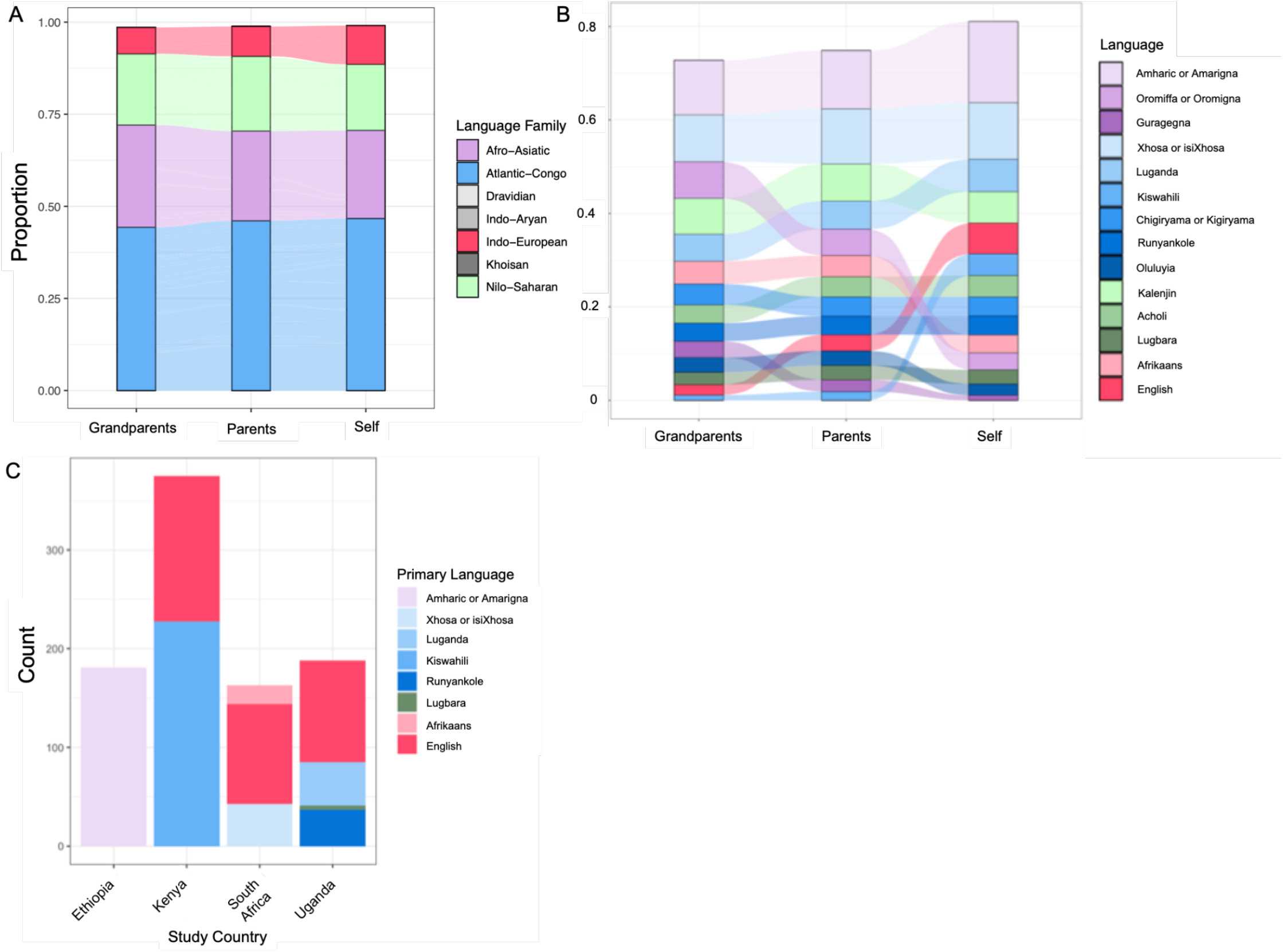
Primary self-reported language shifts over three generations. A) Individual languages were re-classified into broader language families for comparable granularity. B) All languages reported with at least 3% frequency in any generation are shown across the generations. Note the increase in endorsement of English and drop in Oromiffa/Oromigna in the present generation. C) Primary language reported by the individuals within each NeuroGAP study country.

### Genetic Variation Partitions with Language

To assess the correlation between the primary self-reported language and the genetic partitioning that we observed, we conducted Procrustes analyses to measure the correlation between genetic, linguistic, and geographic variation. Procrustes analysis rotates and scales one set of coordinates to minimize the total distance between it and a second set, providing both a metric of similarity between these sets of coordinates and a visualization of their overlap. We use this to compare each individuals’ first three PCs of genetic variation to their geographic locations. We do not have access to the locations where participants’ live or were born, so we use either the locations at which individuals were sampled (study sites) or the centroids of non-English languages that they reported speaking (language based). By including a database of phonemes (units of sound) found in the self-reported languages of individuals and their families and comparing these to the first three PCs of autosomal and X chromosome variation, we found consistent correlations between genetic, linguistic, and geographic variation throughout Africa (**Figure 4**, **Table 1**). We also plotted the genetic PCs, superimposed onto geography via Procrustes, to visualize the geographic distribution of genetic variation (**Supplementary Figure 8**). Because the autosomes and X chromosomes have considerably different numbers of single nucleotide polymorphisms (SNPs), we additionally compared X chromosome variation to chromosome 7, which has a similar length to the X chromosome, and chromosome 22, which is most similar in SNP count to chromosome X (variant counts with/without reference panel intersection: X = 603/1348, chr22 = 705/1455; **Supplementary Figure 9**). To measure linguistic variation, we queried the PHOIBLE 2.0 phonemic database (Moran and McCloy, 2019), which contains phoneme inventories and phoneme qualities for many languages around the world. The resulting matrices of mean phoneme presences were used in a PCA to create four sets of linguistic PCs: a score from languages spoken by the participant, a combined score from those spoken by matrilineal relatives (the participant’s mother and maternal grandmother), a combined score from those of patrilineal relatives (father and paternal grandfather), and a composite score that includes all relatives weighted according to relatedness to the individual (1 for languages spoken by each individual, 0.5 for those of parents, and 0.25 for grandparents). Here, ‘matrilineal’ and ‘patrilineal’ refer to the traits associated with direct lines of descent following exclusively mothers and fathers. We use the languages spoken by these relatives to understand whether sex-biased language transmission may have taken place, and whether it parallels sex-biased gene flow. By comparing linguistic variation associated with these relatives to genetic data and spatial positions, we can explore whether norms and traditions have shaped linguistic and genetic variation.

**Figure 4.**
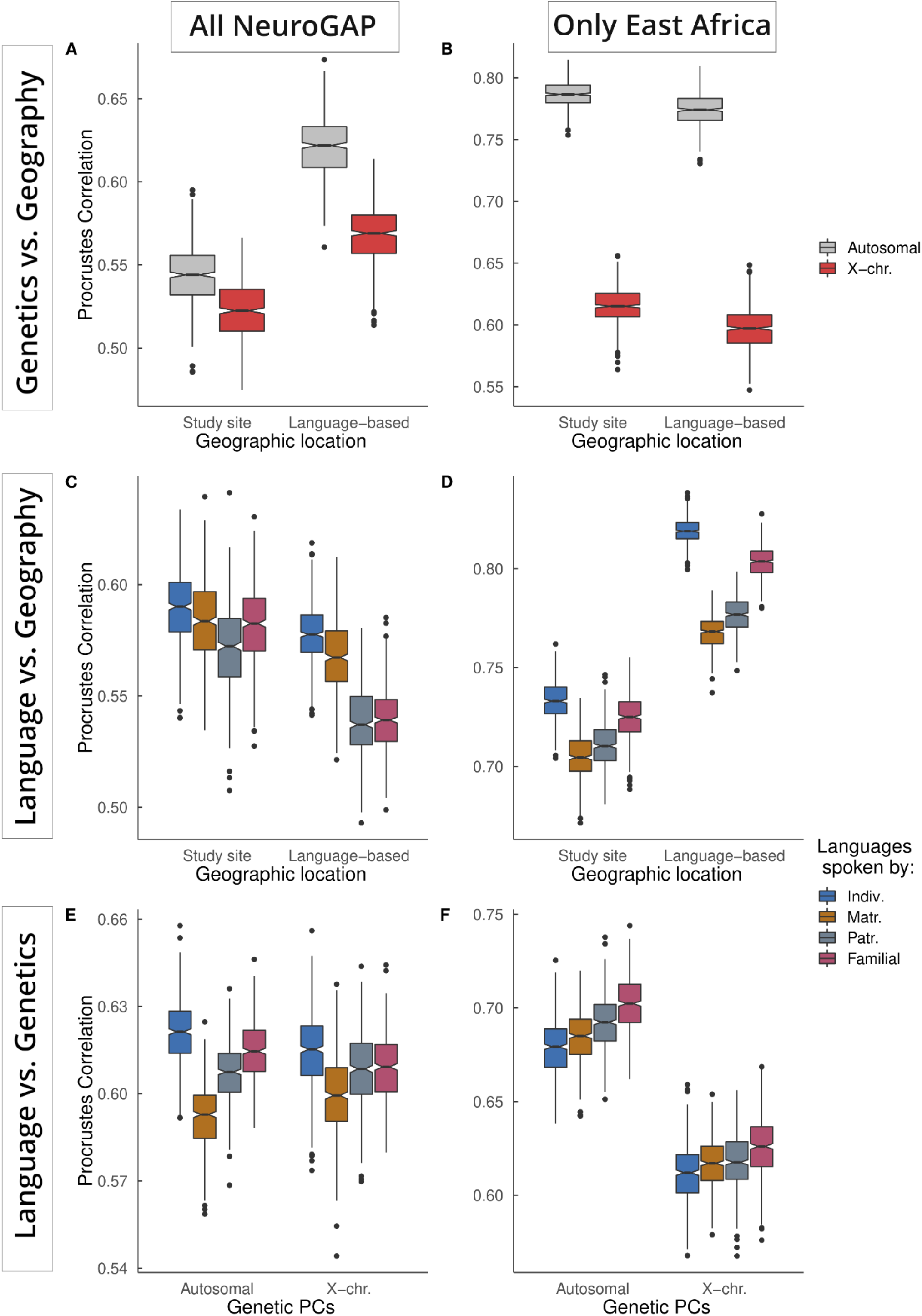
Procrustes correlations between genetics, geography, and language (all p < 5E-5). Procrustes correlations are shown between: **A,B**) geography and genetics. **C,D)** geography and language, and **E,F)** genetics and language. The left column includes results for the entire NeuroGAP collection. The right column contains results subset to the four cohorts in East Africa. For linguistic analyses, linguistic variation is measured by the first three PCs of phoneme inventories from languages reported by individuals as spoken by themselves and their relatives. Matrilineal relatives include the mother and maternal grandmother. Patrilineal relatives include the father and paternal grandfather. Familial refers to a weighted average of all reported family members. Note that Y-axis labels vary between plots.

**Table 1.** Procrustes correlation between genetics, geography, and language. All p < 5E-5. The first three PCs of autosomal and X chromosome variation were used for comparisons. Linguistic variation was calculated as a function of mean phoneme presence across all languages reported by the individual across their pedigree.

| Subset of individuals | PCs 1 to 3:<br>Genetic | Geography | Self | Languages spoken by |  |
| --- | --- | --- | --- | --- | --- |
|  |  |  |  | Mother & Maternal Grandmother | Father & Paternal Grandfather |
| All individuals | Autosomal | 0.5426 | 0.6223 | 0.5935 | 0.6078 |
|  | X chr. | 0.5231 | 0.6078 | 0.5988 | 0.6082 |
| East Africa | Autosome | 0.7868 | 0.6815 | 0.6856 | 0.6924 |
|  | X chr. | 0.6170 | 0.6103 | 0.6178 | 0.6200 |

The first three PCs of both autosomal and X chromosome variation are less correlated to geography (*ρ*=0.543 and 0.533; *p*<5E-5) than are the first three PCs of linguistic variation (*ρ*=0.589; p<5E-5). Both autosomal and X chromosomal variation are similarly correlated to linguistic variation when considering all individuals together. Looking within East Africa alone, X chromosome variation is less correlated to geography (**Table 1**, **Figure 4**). Similarly, linguistic variation associated with individuals’ matrilineal relatives is less correlated to geography and to genetics than that of patrilineal relatives (**Figures 4D, 4F**).

### Language Transmission Through Families

To leverage the detailed multi-generational ethnolinguistic information (see STAR *Methods* “*Ethnolinguistic Phenotypes*”), we computed overall transmission rates of language families over three generations. We initially examined the raw self-reported information of the participant with respect to the primary, second and third language spoken. We assessed the frequency with which the primary language reported by the participant matched each of their older relatives’ (i.e. maternal and paternal grandparents, mother and father) primary language as well as the frequency with which the participants’ primary reported language matched any of the languages reported for their relative (**Table 2**). We find that transmission rates are similar between family members of the same generation when looking at primary language matching any language, regardless of whether including or excluding English.

**Table 2.**
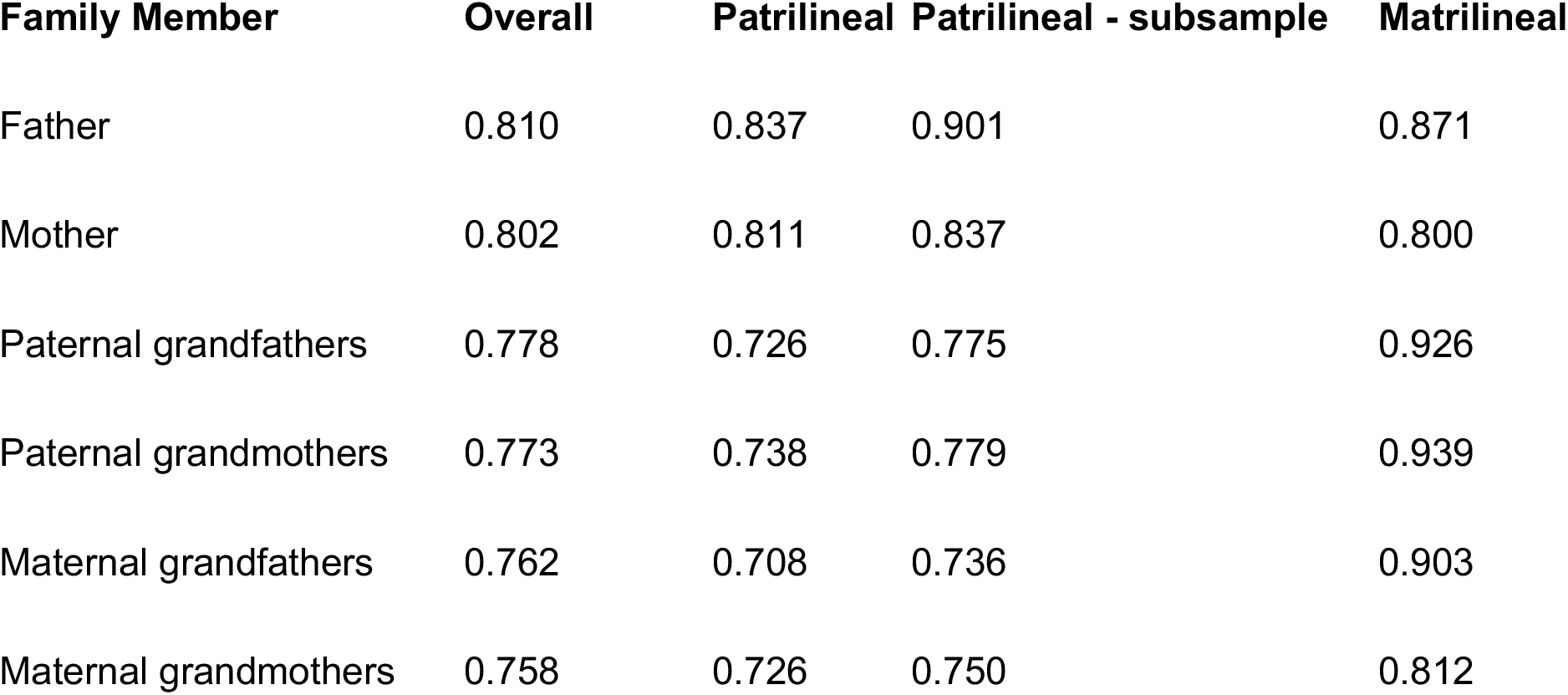
Language transmission rates from relatives. Frequency of a participants’ reported primary language matching one of the top three reported languages spoken by relatives. Rates were calculated excluding English. Given all but one of the NeuroGAP populations with linguistic data were collected in East Africa, we conducted an additional suite of analyses zooming into this region to examine transmission in this part of the continent. In East Africa, individuals were thus additionally partitioned by their affiliation with ethnic groups with either a matrilineal or patrilineal transmission of movable property. Patrilineal languages were run in their entirety as well as subsampled to 105 to match the sample size available for matrilineal languages.

To take a closer look at language transmission across the pedigree, we calculated the rate of transmission between various relatives in our family tree. We note that we only have information for the four countries present in the NeuroGAP dataset, so our results do not capture the full breadth of ethnolinguistic diversity across the continent. In these calculations, we ran tests excluding English to get a better sense of the transmission of languages that have been present in the continent for a longer period of time. We additionally identified whether individuals came from ethnic groups in which the transmission of movable property was historically matrilineal or patrilineal using according to the Ethnographic Atlas (EA)^24^. We then recalculated the transmission rates within those two classifications. Here, we were interested in measuring whether the inheritance of property through the male (patrilineal) or female (matrilineal) lines parallels the transmission of languages. Partitioning east African individuals by the presence of matri- vs patri-lineneal transmission of movable property in their traditional societies (from Murdock’s Ethnographic Atlas, code *EA076(Kirby et al., 2016; Murdock et al., 1999)*), we see a significantly higher transmission rate from individuals assigned to a matrilineal classification (*p*=0.028). As our sample size for matrilineal groups is quite small (N=105 and 674 for matrilineal and patrilineal transmission respectively), we subsampled the patrilineal groupings to the same size as matrilineal and re-ran our analyses. Considered altogether, the trend disappears (*p*=0.097). However, when looking at all familial relationships above the current generation (parents and grandparents) we do detect a significantly higher language transmission rate for individuals assigned to matrilineal groups (*p*=0.012).

The results from our various tests of language transmission and those from Procrustes analysis support the anthropological data that classified the peoples in the regions of Africa that we studied as patrilocal(Vansina 1966; Murdock et al. 1999), but how language was transmitted is uncertain. The lower geography-X chromosome correlation in East Africa suggests that the cultural norms (a predominance of patrilocality) have had effects on genetic variation. We cannot conclusively determine whether language transmission here was historically sex-biased, as recent changes to cultures (including movement to urban areas, colonial histories, access to markets, etc.), has affected the associations between groups of people and languages. There is no evidence of recent sex-baised language transmission in this region from our linguistic data alone. However, the decreased association between X-chromosome variation and language (Fig. 4B) suggests that east Africa has a history of patrilineal language transmission, which parallels the region’s historically predominant patrilocal social structure(Murdock et al., 1999).

Interestingly, we also find that twelve languages reported for earlier generations were not spoken by the participants, indicating that they have disappeared from our dataset. Khoekhoe, Somali, and Urdu disappeared in the parental generation, and Amba, Afar, Argobba, Gumuz, Harar, Hindi, Soddo, Soo, and Tamil were no longer reported languages in the participants’ generation. We caution, however, that many of these languages were observed at very low rates overall.

### Testing for Evidence of Sex-biased Demography

To examine if there was evidence for sex-biased gene flow in our samples, weassessed the similarity of ancestry proportions on the X chromosome versus autosomes. Ancestry fractions were highly correlated across these genomic regions, indicating no evidence for sex-biased demography at this scale, although care should be taken in interpretation given the difference in effective sample size for the X chromosome as compared to the autosomes (**Supplementary Figures 10, 11**). Wilcoxon signed rank tests comparing the fractions of ancestry on X versus autosomes from ADMIXTURE at *k*=4 did not find a significant difference in the means, nor for PC1 vs PC2 (*p* = 0.3754). Similarly, the Mantel tests indicated an observed correlation of 0.987 (simulated p-value= 0.001) for the X-chromosome compared to the autosomal F_ST_ values. We additionally ran Procrustes analyses comparing genetic and linguistic variation on the X chromosome as compared to the autosomes in their entirety, chromosome 7 - most similar in length to chrX - and chromosome 22 -most similar in SNP count to chrX (Supplementary Figure 9). Similarly, the Procrustes tests showed significant correlation between the first three PCs of X and autosomal variation (*ρ*=0.850 for all of Africa and *ρ*=0.753 for East Africa alone). Compared to chromosomes 7 and 22,, results were similar (*ρ*=0.826 and 0.780 for all Africa, and *ρ*=0.728 and 0.691 for East Africa).

### Reference Panels Miss Meaningful Allele Frequency Resolution within Africa

We visualized allele frequencies for functionally important variants across our 5 collection sites as compared to reference data from the 1000 Genomes Project. One example variant (rs20713348), key in beta-thalassemia, varies in frequency depending on the precise location in Africa considered. The NeuroGAP-Psychosis allele frequencies observed across the five sites were: Ethiopia:12.26%, Kenya (KEMRI):11.35%, Kenya (Moi): 9.71%, Uganda:13.11%, and SouthAfrica:5.7%. As this variant has direct consequences on human health, consideration of the difference in frequency across the continent is meaningful. For another example, rs72629486 (**Supplementary Figure 12**), a missense coding single nucleotide variant in the gene *ACTRT2*, ranges in minor allele frequency (MAF) in NeuroGAP-Psychosis from 5% in Ethiopia down to 1.3% in Uganda (other frequencies were: Kenya (KEMRI): 2.6%, Kenya (Moi): 2.6% and South Africa:2.3%). This is nearly the full range of the frequency distribution for all global populations in the gnomAD database(Karczewski et al., 2020), which lists the variant in the AFR as 5.5%, missing finer subcontinental resolution. rs72629486 is predicted to be deleterious and probably damaging by SIFT and PolyPhen, respectively, and has a combined annotation dependent depletion score of 22.9, highlighting that this variant is likely to be highly functionally important(Adzhubei et al., 2013; Ng and Henikoff, 2003; Rentzsch et al., 2019).

## Discussion

Africa is a highly diverse continent, home to immense genetic, linguistic, and cultural diversity. This ethnolinguistic variation is extremely complex and is meaningful to disentangle prior to statistical genetics analyses. Here, we measured the correlation between genetic, linguistic, and geographic variation focusing on four African countries for which we have collected data as part of the NeuroGAP-Psychosis study. We find that genetic and linguistic variation are closely correlated to each other as well as to geography across the African continent. This is consistent with previous work examining global patterns of diversity as well as the ‘Bantu expansion’, one of the largest demographic events in African history(Baker et al., 2017; Beleza et al., 2005; Creanza et al., 2015; de Filippo et al., 2012; Li et al., 2014; Uren et al., 2016). We find that across the regions of Africa that we surveyed, language is closely correlated to both genetics and geography, a phenomenon that has been noted in Europe and Ethiopia previously(Longobardi et al. 2015; Piazza et al. 1995; Pagani et al. 2012; López et al. 2021; Barbujani et al. 1994). The patterns of linguistic and genetic variation in this dataset suggest a history of patrilocal residence, in accord with previous studies of the region(Wood et al. 2005; Destro-Bisol et al. 2004; Vansina 1966). We find here that individuals collected from the same geographic location show significant genetic differentiation by language family, particularly in East Africa, where there is immense linguistic diversity. Previous studies have examined this covariation between culture and genetics produced by a history of migrations and population expansions(Tishkoff et al. 2007; Hollfelder et al. 2017; Gomes et al. 2015; Pagani et al. 2012), and we explore how this affects genetic datasets by examining both genetics and ongoing changes to culture. These data should be synthesized to influence how population substructure is controlled for in genetic tests, and we suggest that anthropological data should be incorporated into a nuanced treatment of genetic clusters. For example, future work exploring the direct incorporation of ethnolinguistic affiliations into linear mixed models would be useful, e.g. in the context of including it as a random effect covariance matrix to better control for stratification(Heckerman et al., 2016).

As there is such immense genetic variation across the African continent(1000 Genomes Project Consortium et al., 2010; Bergström et al., 2020; Choudhury et al., 2020; Gurdasani et al., 2015, 2019), we highlight cases where such variability may be particularly informative. Africa is not simply one monolithic location, as it is sometimes treated in major genomics resources that include primarily or exclusively African Americans as representatives of genetic variation for the entire African continent(Karczewski et al., 2020; Taliun et al., 2021). Rather, there is an extraordinary amount of genetic variability within it. These example loci highlight both the diversity of variation within the African continent, as well as the fact that within-Africa variation can be informative for broader variant interpretation; many variants appearing rare elsewhere are common in parts of Africa.

As part of the NeuroGAP-Psychosis study’s recruitment process, multi-generational self-reported ethnolinguistic data was collected from participants, including individual ethnicity and at least primary, second and third language from participants for themselves, as well as for each of their parents and grandparents. This provides us with an unusually rich depth of multigenerational demographic information from participants, a unique strength of our dataset that affords us the opportunity to investigate language transmission through the pedigree and shifts in language frequencies over time. First, we examined the overall change in self-reported language frequencies over three generations. Perhaps most striking is the increase in the reporting frequency of English by participants as their primary language as compared to their reports for older generations of their family. Twelve languages reported for earlier generations were no longer reported as spoken by the participants, suggesting their loss from this cohort. Interestingly, these languages represent a mix of both historically spoken and imported languages for the countries that enrolled participants in the NeuroGAP-Psychosis study. These results emphasize that the analysis of phenotypes should consider not only how they relate to genetics, but how phenotypes may be affected by a rapidly changing cultural environment. While these results are intriguing, we stress that our participants are not necessarily representative of the local populations from which they come and do not begin to cover the full breadth of variation across the African continent. A further consideration is a potential upwards bias towards reporting of English and Amharic as a primary language due to a preference towards reporting the language of consent as primary, as well as towards languages taught in local educational systems. This additionally highlights the importance of careful consideration of items on self-report forms to ensure accurate and representative phenotype collection.

In summary, better understanding the composition of samples is a key first step to calibrating subsequent statistical genetics analyses. Cultural factors such as language can dramatically impact the structure of cohort data; we find that self-reported language classifications meaningfully track underlying genetic variation that varies independently from geography. The work presented here improves the understanding of the immense spectrum of genetic and ethnolinguistic variation found across multiple African populations and sheds light on the shifts in reported language over the past three generations in five collection sites.

## STAR Methods

### Collection Strategy

As described in more detail in the published protocol(Stevenson et al., 2019), NeuroGAP-Psychosis was designed as a case-control study recruiting participants from more than two dozen hospitals and medical clinics in Ethiopia, Kenya, South Africa, and Uganda. Participants were recruited in languages in which they are fluent, including Acholi, Afrikaans, Amharic, English, Kiswahili, Luganda, Lugbara, Oromiffa/Oromigna, Runyankole, and isiXhosa. After consenting to be in the study, participants gave a saliva sample using an Oragene kit (OG-500.005) for DNA extraction. Study staff then asked a range of questions on demographics, mental health, and physical health and took participants’ blood pressure, heart rate, height, and weight. The whole study visit lasts approximately 60-90 minutes. **Supplementary Table 1** contains details about the dataset and country of origin of all populations included in analyses in this manuscript.

### Ethnolinguistic Phenotypes

Multiple phenotypes related to self-reported ethnolinguistic categorizations have been collected as part of the recruitment process. This includes multi-generational data including each participants’ birth country as well as primary, secondary and tertiary language and ethnicity. All linguistic data were collected from participants both for themselves as well as for each of their parents and grandparents, giving an unusually rich depth of information. The specific phrasing of questions collected are as follows:

Primary language (lang_self_1): “What primary language do you speak?”
2nd language (lang_self_2): “What 2nd language do you speak?”
3rd language (lang_self_3): “What 3rd language do you speak?”
Primary-3rd ethnicity (ethnicity_1): “What is your ethnicity or tribe?”

Reports for other relatives followed similar phrasing. The primary language question for each is listed, with primary swapped for ‘2nd’ or ‘3rd’ for the second and third reported languages for that family member.

Mother (lang_mat_1): “What was the primary language that your biological mother spoke?”
Father (lang_pat_1): “What was the primary language that your biological father spoke?”
Maternal grandmother (lang_mgm_1): “What primary language did your biological mother’s mother speak?”
Maternal grandfather (lang_mgf_1): “What primary language did your biological mother’s father speak?”
Paternal grandmother (lang_pgm_1): “What primary language did your biological father’s mother speak?
Paternal grandfather (lang_pgf_1): “What primary language did your biological father’s father speak?

**Supplementary Table 2** contains raw data for language transmission counts for all languages reported in the NeuroGAP-Psychosis dataset. **Supplementary Table 3** indicates matrilineal or patrilineal classification of all self-reported ethnicities in the dataset.

### Genetic Data Quality Control

Quality control (QC) procedures for NeuroGAP-Psychosis data were done using the Hail python library (www.Hail.is). All of the data was stored on Google Cloud. The QC steps and filters used were adapted from Ricopili(Lam et al., 2019) and Anderson et al. 2011(Anderson et al., 2010). The data was genotyped using the Illumina Global Screening Array. For each of the five NeuroGAP-Psychosis sites, a VCF file with genotyping data was stored on Google Cloud. Before QC, each VCF contained 192 samples and 687537 variants. When looking at the data pre-QC we discovered elevated deviations in Hardy Weinberg Equilibrium. We found that autocall call rate, Illumina’s custom genotype calling algorithm, explained these deviations. The QC filtering steps thus took place after removing individuals with an autocall call rate less than .95. 937 of the original 960 individuals remained. These 960 individuals were used for the linguistic transmission analyses presented here (with some missing data for specific familial categories), while for genetic analyses further QC on variants was conducted.

The site-specific VCF files were imported as Hail matrix tables and annotated with appropriate data from the metadata file before being merged. The resulting matrix table had 937 samples and 687537 variants. Prior to QC, the joint dataset was split into autosomes, PAR, and nonPAR regions of the X chromosome. QC filtering was conducted separately for the autosome and X chromosome regions. Pre-QC, the autosomal dataset had 937 samples and 669346 variants. The following is a list of the QC steps and parameters used for autosomal QC. (1) Removing variants with a call rate less than 95%. After filtering, 638235 variants remained. (2) Removing individuals with a call rate less than 98%. After filtering, 930 individuals remained. (3) Removing individuals whose reported sex did not match their genotypic sex. After filtering, 923 individuals remained. (4) Removing variants with a minor allele frequency less than 0.5%. After filtering, 360,321 variants remained. This large drop in variants was expected as the GSA array poorly tags common variation in samples with African Ancestry(Martin et al., 2021). (5) Removing variants with a Hardy Weinberg Equilibrium *p*-value less than 1 × 10^−3^. After filtering, 331667 variants remained. (6) Using PC-Relate with 10 PCs, removing individuals with a kinship coefficient greater than .125. After filtering, 900 individuals remained. After autosomal QC, 900 individuals and 331667 variants remained.

The PAR and nonPAR regions of the X chromosome were subset to the 900 samples which passed autosomal QC before going through variant QC. The same variant thresholds used for autosomal QC were used to conduct QC on the PAR region. Pre-QC, the PAR region had 900 samples and 518 variants. (1) After SNP call rate filtering, 515 variants remained. (2) After MAF filtering, 411 variants remained. (3) After HWE filtering, 402 variants remain. Post-QC, the PAR region had 900 samples and 402 variants. For the nonPAR region, the dataset was split by sex. The female nonPAR dataset had 441 samples and 17673 variants. Variant QC was carried out on the females using the following metrics. (1) Removing variants with a call rate less than 98%. After filtering, 16261 variants remained. (2) Removing variants with a minor allele frequency less than 1%. After filtering, 11113 variants remained. (3) Removing variants with a Hardy Weinberg Equilibrium *p*-value less than 1 × 10^−6^. After filtering, 11104 variants remained. After nonPAR QC on the females, the male nonPAR dataset was merged with the female QC’d nonPAR dataset. The final nonPAR dataset had 900 samples and 11104 variants. After QC, the autosomal, PAR, and nonPAR datasets were merged into one matrix table.The final merged post-QC dataset contained 900 samples and 343173 variants. The counts of variants/individuals per site after autosomal and X QC can be found in **Supplementary Tables 4 and 5**.

After QC, the dataset was merged with three reference panel datasets, the 1000 Genomes Project (1kGP(Auton and Salcedo 2015; Byrska-Bishop et al. 2021)(Auton and Salcedo, 2015) and the African Genome Variation Project(AGVP)(Gurdasani et al., 2015). Before merging with HGDP and 1kGP, the AGVP and NeuroGAP datasets were merged and lifted over from GRCh37 to GRCh38. Before this initial merge,, AGVP had 1297 samples and 1778578 variants while NeuroGAP had 900 Samples and 343173 variants. Prior to merging these two datasets, multi-allelic variants were removed from the NeuroGAP dataset resulting in 343165 variants. The two datasets were then combined using plink −bmerge. The resulting dataset had 2197 samples and 1908204 variants after intersection. A 5% geno filter using the −geno plink command was then run on the dataset which gave the final merged dataset counts prior to liftover of 2197 samples and 206240 variants. The liftover of the merged AGVP, NeuroGAP dataset was conducted using Hail. After liftover, there were 20197 samples and 206156 variants. Next, the AGVP NeuroGAP dataset was merged with a subset of the gnomAD v3.1 release which consisted of newly sequenced HGDP and 1kGP datasets<u>(GnomAD v3.1 release blog)</u>. This dataset of HGDP and 1kGP had 4097 samples and 155648020 variants prior to merging. After combining AGVP plus NeuroGAP with HGDP plus 1kGP using plink −bmerge, the resulting dataset contained 6294 samples and 149518 variants. A 5% geno filter was then run on the dataset which resulted in the final counts of 6294 samples and 148488 variants for the NeuroGAP and reference panels dataset.

### Population Structure and Admixture Analyses

Cohort data from the five NeuroGAP-Psychosis plates were merged with African reference populations from the 1000 Genomes Project(Byrska-Bishop et al. 2021; Auton and Salcedo 2015) and the African Genome Variation Project(Gurdasani et al., 2015). These populations provide reasonably comprehensive geographic coverage across the African continent from currently available reference panels and contain populations which are co-located in the same countries as all NeuroGAP-Psychosis samples. We note, however, that there are additional datasets of African genetic variation beyond the reference panels incorporated into these analyses, including H3Africa, the Uganda Genome Resource, and many cohorts from specific collection sites(Pagani et al. 2015; Gurdasani et al. 2019; Choudhury et al. 2020). PCA was run using flashPCA(Abraham et al., 2017). Detailed examination of admixture was conducted using the program ADMIXTURE(Alexander et al., 2009) with five-fold cross validation error to inform the correct number of clusters. We used the unsupervised mode of ADMIXTURE ten times for each value of k to capture any different modes present in the data. All runs treated data as diploid. Plots from ADMIXTURE output were generated with pong(Behr et al., 2016). ADMIXTURE was run using a tailored representation of global genetic data consisting of all continental African populations, the CHB population from China to capture East Asian admixture, the GBR from Britain to capture European admixture, and the GIH from India to capture South Asian ancestry. Regions were assigned as according to the UN Statistics division geoscheme<u>(United Nations Statistics Division)</u>.

F_ST_ estimates across populations were generated using smartPCA(Patterson et al., 2006). F_ST_ heatmaps were generated in R using the package *corrplot* (Wei and Simko 2017). The relationship between ancestry composition on the autosomes vs X chromosome was examined using Wilcoxon rank tests and Mantel tests in R with the package *ade4* (Jombart 2008).

### Relationship between Genetics and Language

To measure linguistic variation, we made use of the PHOIBLE 2.0 phonemic database(Moran and McCloy, 2019), which contains phoneme inventories and phoneme qualities for languages around the world. For every individual, we identified all languages spoken—excluding English—that were present in the PHOIBLE database (84.5% of languages spoken by the individuals themselves, and 81.1% of languages spoken by their relatives). Using the phoneme inventories (including both primary phonemes and their allophones) from PHOIBLE, we found the mean phoneme presence for each individual’s or each relative’s spoken languages (if one of two spoken languages contained the sound /g/, the /g/ value for that individual would be 0.5). The resulting matrices (of individuals or their relatives, and mean phoneme presences) were transformed using PCA conducted in R to create three sets of principal components (PCs): from personally spoken languages, from those spoken by matrilineal relatives (mother and maternal grandmother), and from those of patrilineal relatives (father and patrilineal grandfather).

To observe the broader linguistic changes taking place, all languages were assigned the highest-level classifications available in Glottolog 4.2.1(Hammarström et al., 2020). These classifications were modified to minimize the number of high-level classifications while maintaining an element of geographic origin. Several classifications were consolidated into Nilo-Saharan (made up of Nilotic, Central Sudanic, Kuliak and Gamuz classifications) and Khoisan (Khoe-Kwadi, Kxa, and Tuu), and Afro-Asiatic was expanded (with Ta-Ne-Omotic and Dizoid). Indo-European was split to account for the recent history of its speakers: Afrikaans and Oorlams were placed into a unique category, the languages of Europe into another, and those of the Indian subcontinent (Hindi and Urdu) into a third. We excluded languages that were unclassified or identified as speech registers.

Every individual was associated with a survey location, meaning the geographic coordinates where the sample was collected, and we used the spoken languages to assign a different, linguistic location. To do this, using all languages an individual spoke, and these languages’ locations from Glottolog, we calculated the mean location of each individual’s languages.

To compare linguistic, genetic, and geographic variation, we used a set of Procrustes analyses implemented in R(Dixon, 2003). For linguistic and genetic variation, the first three PCs of variation were used. Since Procrustes minimizes the sum of squared euclidean distances, the geographic coordinates of each individual were converted to points on a sphere. To measure the correlation between geographic variation and linguistic or genetic variation, the linguistic and genetic PCs were transformed (via rotation and scaling) to minimize the sum of squared distance between individuals’ geographic locations and the transformed genetic or linguistic PCs. The first three PCs of Procrustes-transformed linguistic and genetic variation—representing their similarity to geographic variation—were then plotted onto a map. The vegan::protest function was used for permutation testing of Procrustes tests to produce an empirical *p* value threshold for significance. It did 19,999 permutations of the data and all had a lower correlation than we observed, so the empirical *p* value is 1/(19,999 + 1) = 5E-5, as presented in Figure 4.

We additionally calculated the transmission frequency of languages from sets of family members. Given the discrepancy in number of languages in the matri- vs patri-groupings, patrilineal languages were additionally subsampled to the same overall number as matrilineal and rates were recalculated.

### Anthropological variables

To identify relevant anthropological data, we accessed data from the Ethnographic Atlas (EA)(Murdock et al., 1999) using D-PLACE (Kirby et al., 2016). We associated each ethnicity reported in the NeuroGAP-Psychosis survey data to a society in the EA (if possible), and used variable *EA076: Inheritance rule for movable property*. For ethnicities with data, individuals whose ethnicities were associated with consistent inheritance rules or marital residence patterns were assigned that rule or pattern. Of the 907 NeuroGAP-Psychosis individuals 779 were assigned an inheritance rule (matrilineal or patrilineal). Additionally, 751 individuals could be assigned a post-marital residence rule (patrilocal, neolocal, or virilocal-like) using *EA012: Marital residence with kin*, but matrilocality and other forms of residence were not found among the sampled ethnicities, and we did not use these for our analyses. Similarly, other variables such as *EA074: Inheritance rule for real property (land*) did not vary for available individuals (all available ethnicities traditionally practiced either patrilineal, male-biased, or neutral patterns of real property inheritance). Only a single ethnicity corresponding to eight individuals could be assigned a matrilineal inheritance pattern based on *EA043:Descent: major type*.

## Acknowledgements

We thank all participants for their willingness to contribute their data to this effort. John Mugane and Ana Maria Olivares provided advice and assistance related to this project. This work was supported by funding from the NIH/National Institute of Mental Health (K01MH121659 and T32MH017119 to E.G.A.; K99/R00MH117229 to A.R.M.; L.B.C., K.C.K, B.G., D.J.S, S.T., and D.A. are supported in part by R01MH120642).

## Author Contributions

Conceptualization, E.GA.; A.R.M.; K.C.K.; Formal Analysis, E.G.A., S.D., Y.P., A.K., L.M.; Data Curation, M.B., Z.K., C.P.N., A.S., R.E.S; Investigation: S.G., J.K., R.J., R.M.M., M.A., W.E.I., G.K., W.S., F.K.A., H.M., L.M.; Resources, T.A., D.A., M.A., F.K.A., L.A., A.F., S.G., W.E.I., R.J., S.M.K., G.K., E.K., J.K., H.M., R.M.M., C.R.J.N., R.R., W.S., D.J.S., S.T., Z.Z.; Writing - Original Draft, E.G.A., A.R.M.; Writing - Review & Editing, all authors; Visualization, E.G.A., S.D., Y.P., A.K., L.M., Z.K.; Supervision, N.C., M.J.D, B.M.N., K.C.K., S.R., A.S., T.A., D.A., L.A., A.F., W.E.I., S.M.K., G.K., E.K., C.P.N., C.R.J.N., R.R., D.J.S., R.E.S., S.T., Z.Z., L.B.C., B.G.; Project Administration, A.S., R.E.S., M.A., S.G., R.J., J.K., L.O., R.M.M., C.P.N.; Funding Acquisition, M.J.D, B.M.N., K.C.K.

## Human Subjects Approval

Ethical clearances to conduct this study have been obtained from all participating sites, including:

· Ethiopia: Addis Ababa University College of Health Sciences (#014/17/Psy) and the Ministry of Science and Technology National Research Ethics Review Committee (#3.10/14/2018).
· Kenya: Moi University School of Medicine Institutional Research and Ethics Committee (#IREC/2016/145, approval number: IREC 1727), Kenya National Council of Science and Technology (#NACOSTI/P/17/56302/19576), KEMRI Centre Scientific Committee (CSC# KEMRI/CGMRC/CSC/070/2016), KEMRI Scientific and Ethics Review Unit (SERU# KEMRI/SERU/CGMR-C/070/3575)
· South Africa: The University of Cape Town Human Research Ethics Committee (#466/2016)
· Uganda: The Makerere University School of Medicine Research and Ethics Committee (SOMREC #REC REF 2016-057) and the Uganda National Council for Science and Technology (UNCST #HS14ES)
· USA: The Harvard T.H. Chan School of Public Health (#IRB17-0822)

## Data and Code Availability Statement

The genetic data generated during this study for NeuroGAP-Psychosis samples are available on dbGAP (accession number phs2528.v1). Code used to process and analyze data is freely available on Github at: https://github.com/atgu/NeuroGAP.

## Declaration of Interests

A.R.M. has consulted for 23andMe and Illumina and received speaker fees from Genentech, Pfizer, and Illumina. B.M.N. is a member of the Deep Genomics Scientific Advisory Board. He also serves as a consultant for the Camp4 Therapeutics Corporation, Takeda Pharmaceutical and Biogen. M.J.D. is a founder of Maze Therapeutics. The remaining authors declare no competing interests.

## Notes

### Summary of Updates

Updated PDF to the most recent version which has been resubmitted after first round peer review.

