## Supplementary Information for "Genetic structure correlates with ethnolinguistic diversity in eastern and southern Africa"

*Finer scale description of genetic structure across NeuroGAP-Psychosis countries*

*Ethiopia*: The pilot data from Addis Ababa University (AAU) falls cleanly within the Ethiopian reference panel cluster, as would be expected by the collection location in Addis. This also matched with the fact that the majority of the participants' self-reported languages were Amhara and Oromo, and we have reference panels from these corresponding ethnic groups from the AGVP. Individuals from Ethiopia tend to be quite genetically distinct from people from other areas of Africa, pulling out a unique ancestral component at K=4, immediately after the separation of European and east Asian individuals from Africa. They also appear to have some European admixture, visible as the red component in ADMIXTURE plots (Figure 1A). This may be related to back-migration into the continent (López et al., 2021; Pagani et al., 2012, 2015; Pickrell et al., 2014).

*Kenya*: The pilot data from Moi University falls within the East African cluster, as would be expected by the collection location in Eldoret (Figure 1B). Furthermore, it seems to fall with the Kalenjin and Luhya ("LWK") groups primarily, which are the most common self-reported ancestry that participants reported in these 192 samples (Figure 1). Interestingly, two geographically close East African populations (shown in red) dispersed into distinct clusters, which by PC5 define that axis of variation. We next investigated features that might explain this differentiation between closely geographically oriented groups. The two distinct red East African groups appear to speak different languages, one speaking an Afro-Asiatic language and one a Niger-Congo, such they function as reasonably independent groups genetically even though they are in very close geographic proximity to one another.

The pilot data from the KEMRI-Wellcome Trust also overlaps roughly with the East African reference panels, but the core of the pilot samples do not lie squarely on the reference panels. There are a couple of reasons why this might be happening: 1) the reference panels for Kenya are from the Kalenjin and Luhya ("LWK") groups, which are from western Kenya and geographically far away from Kilifi where the participants were recruited. 2) Due to the history of coastal Kenya, there is likely a lot of admixture between people who originated from the coast and people of Arabic ancestry. Admixture is when two historically separate groups of people mix with each other. Unfortunately, there are no reference panels from East African coastal populations or from the Arabian Peninsula. 3) There could be a technical error with the data.

*Uganda*: The pilot data from Uganda also falls cleanly within the East African cluster, as would be expected by the collection location in Kampala. Furthermore, it seems to fall with the Baganda ethnic group primarily, which is the most common self-reported ancestry that participants reported in these 192 samples. We note the breakdown of Ugandan samples by language group in a similar fashion to what we observed in Kenya, and have included them in more detailed analyses of the correlation between genetic similarity and language family divergence.

*South Africa*: The pilot data from the University of Cape Town (UCT) falls most closely to the South African reference panels (in purple) on PC space. However, the core of the pilot samples do not lie squarely on the reference panels. There are several possible explanations for this: 1) the reference panels for South Africa are from the Zulu and the Sotho groups, which are in eastern South Africa and geographically far away from Cape Town and other locations, where the participants were recruited. 2) Cape Town is inhabited by people all over Africa and the world and there are many immigrants living there. Since NeuroGAP-Psychosis does not exclude participants based on ancestry or where they were born, there are likely to be people who were born outside of South Africa taking part in the study, leading to several individuals falling in other geographic areas of Africa. 3) Due to the history of South Africa, with immigration from East Africa, Europe, Malaysia, among other places, and with intermarriage with the indigenous Khoi and San groups, there is a lot of admixture in the Western Cape (Bergström et al., 2020; Chimusa et al., 2015; Pickrell et al., 2014; Sikora et al., 2011; Uren et al., 2016). Indeed, we see indications of admixture in our NeuroGAP-Psychosis UCT samples, both within different African continental groups as well as contributions from other continental groups.

### Supplementary Figures

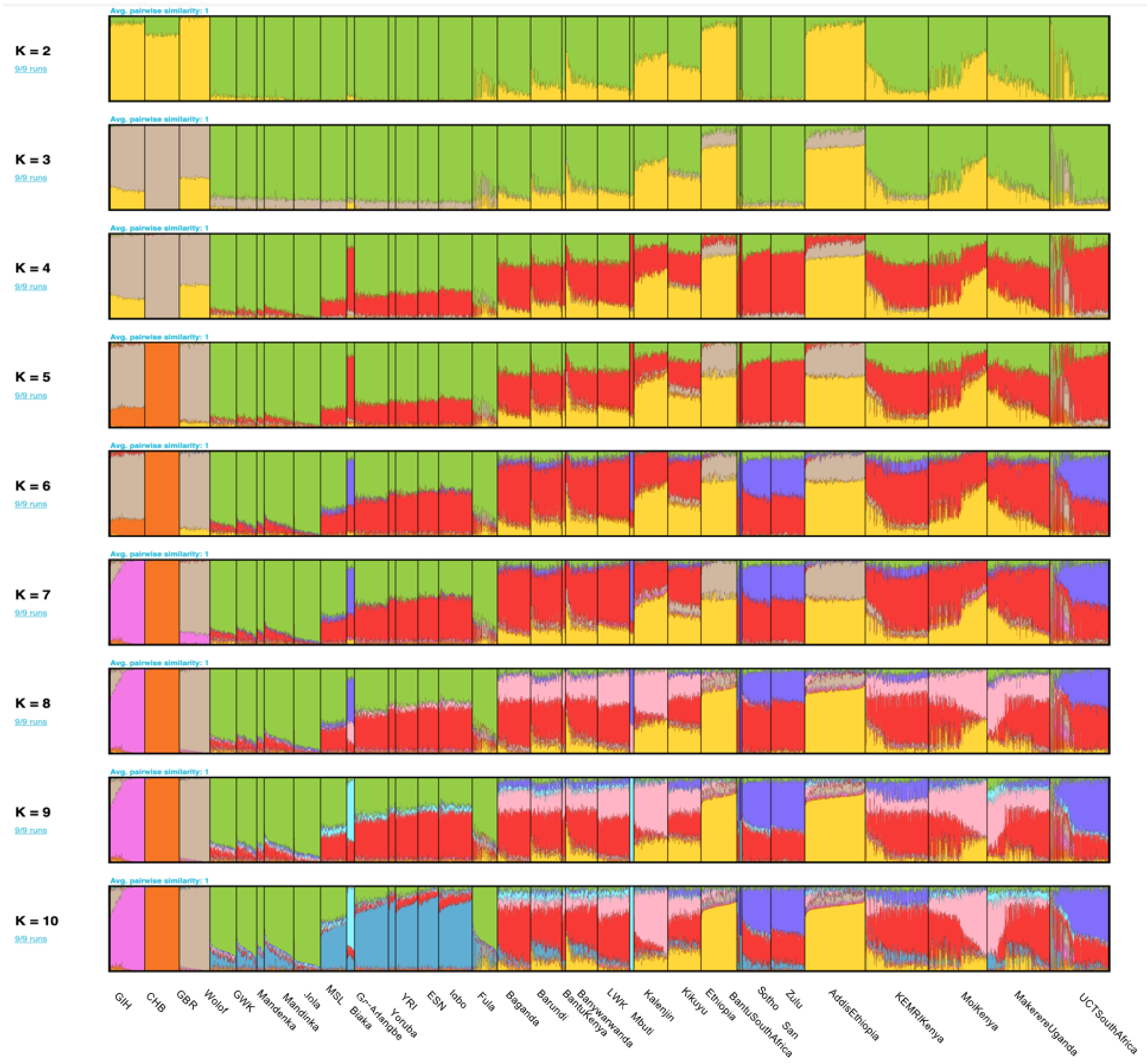

***Supplementary Figure 1.*** *ADMIXTURE plot showing k=2 through k=10 for all African populations as well as a tailored non-African reference panel comprising representation from a south Asian (GIH, fuschia), east Asian (CHB, orange), and European (GBR, tan). A full description of populations, their source datasets, their geographic locations, and their linguistic assignments can be found in Supplementary Table 1.*

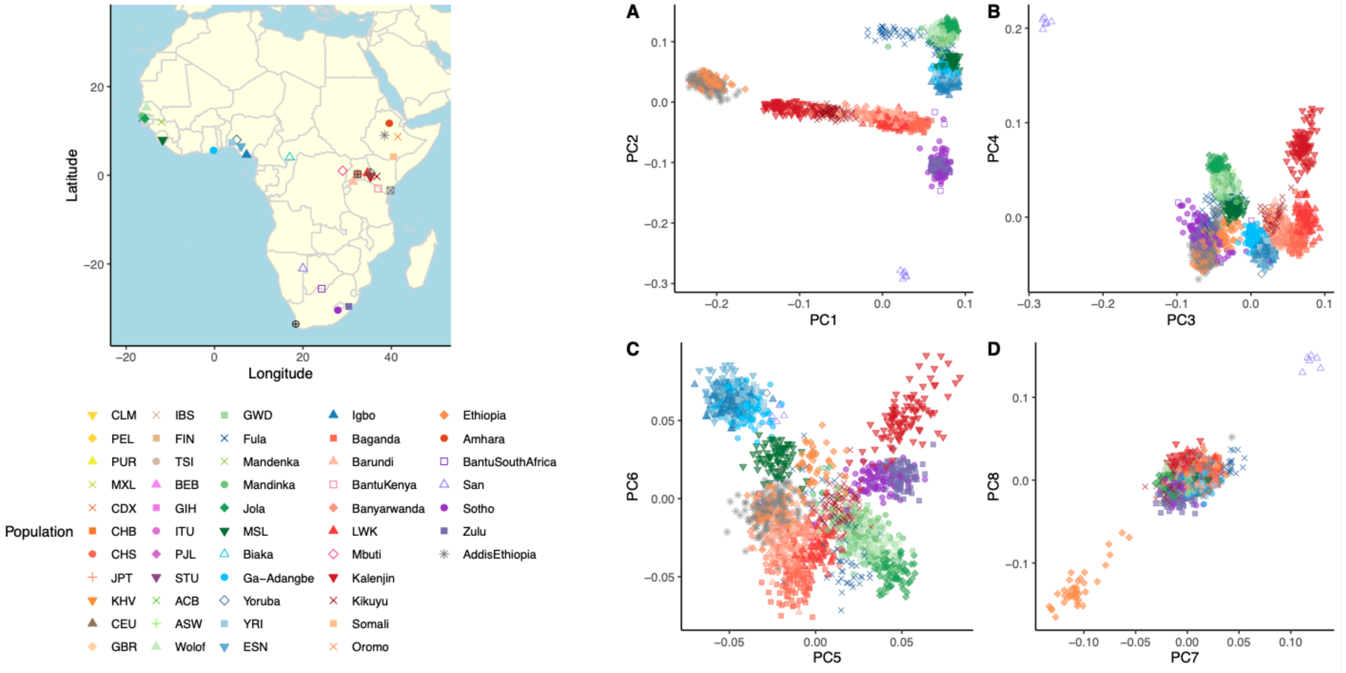


***Supplementary Figure 2****. Fine-scale structure of genetic variation in East Africa. A map showing the location of populations plotted is shown on the left.* ***A-D****) PCA plots for PCs 1-8 showing clustering of AddisEthiopia NeuroGAP-Psychosis samples across PC space with an African reference panel.*

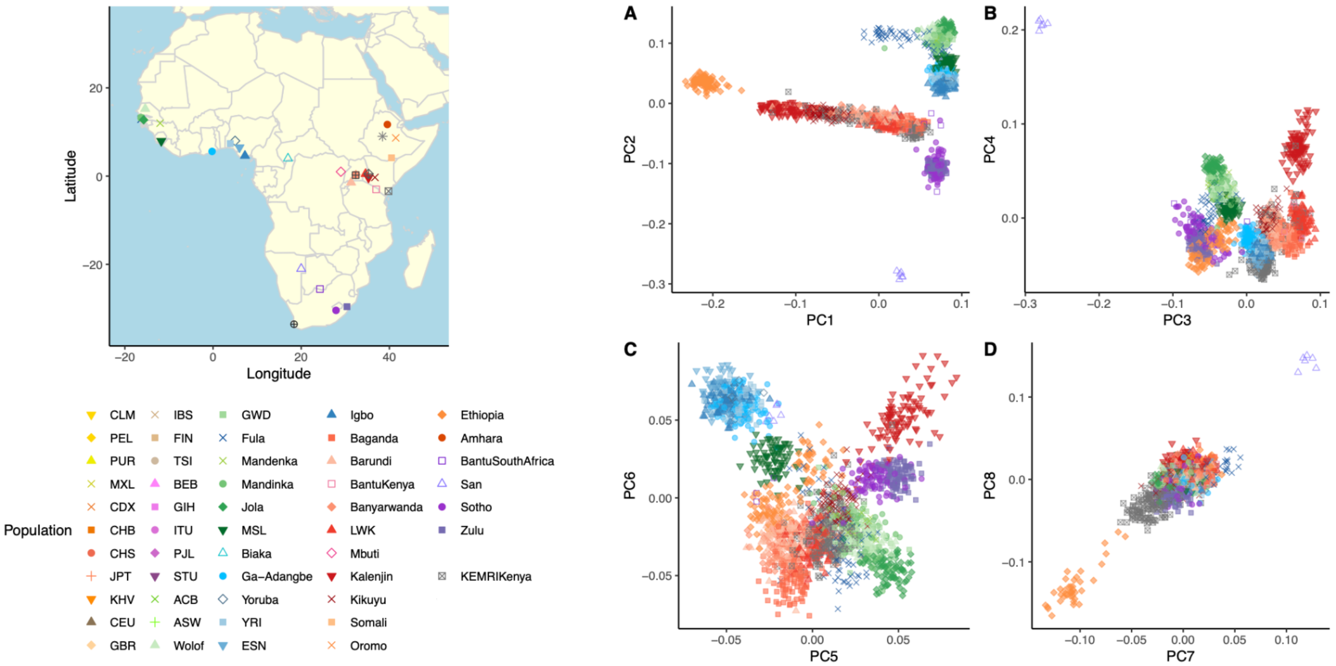


***Supplementary Figure 3****. Fine-scale structure of genetic variation in the KEMRIKenya NeuroGAP-Psychosis collection site. A map showing the location of populations plotted is shown on the left.* ***A-D****) PCA plots for PCs 1-8 showing clustering of KEMRIKenya NeuroGAP-Psychosis samples across PC space with an African reference panel.*

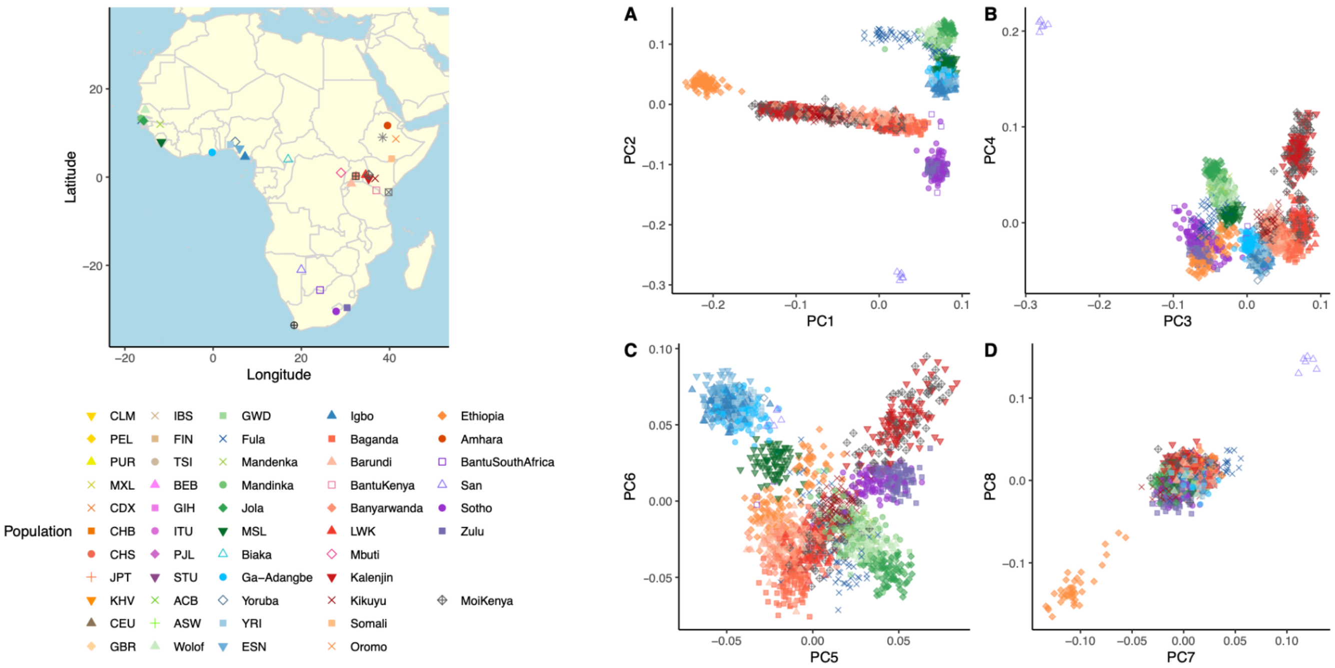


***Supplementary Figure 4***. *Fine-scale structure of genetic variation in the MoiKenya NeuroGAP-Psychosis collection site. A map showing the location of populations plotted is shown on the left.* ***A-D****) PCA plots for PCs 1-8 showing clustering of MoiKenya NeuroGAP-Psychosis samples across PC space with an African reference panel.*

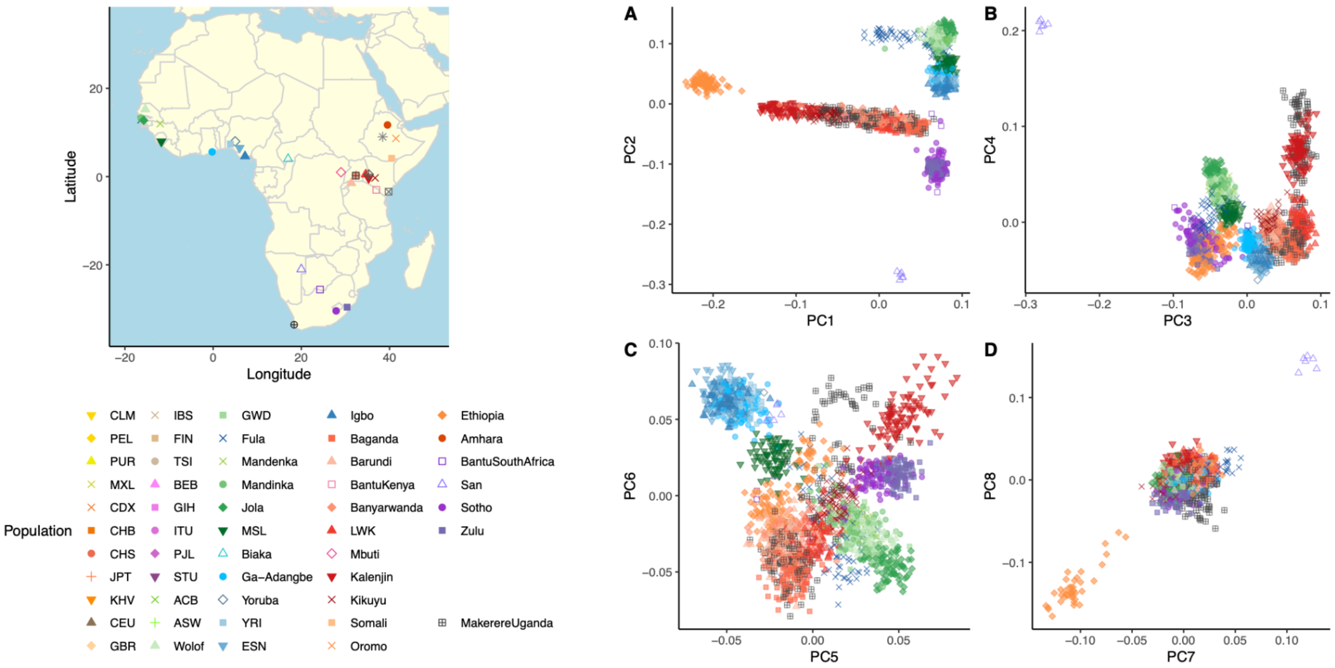


***Supplementary Figure 5**. Fine-scale structure of genetic variation in the MakerereUganda NeuroGAP-Psychosis collection site. A map showing the location of populations plotted is shown on the left. **A-D**) PCA plots for PCs 1-8 showing clustering of MakerereUganda NeuroGAP-Psychosis samples across PC space with an African reference panel.*

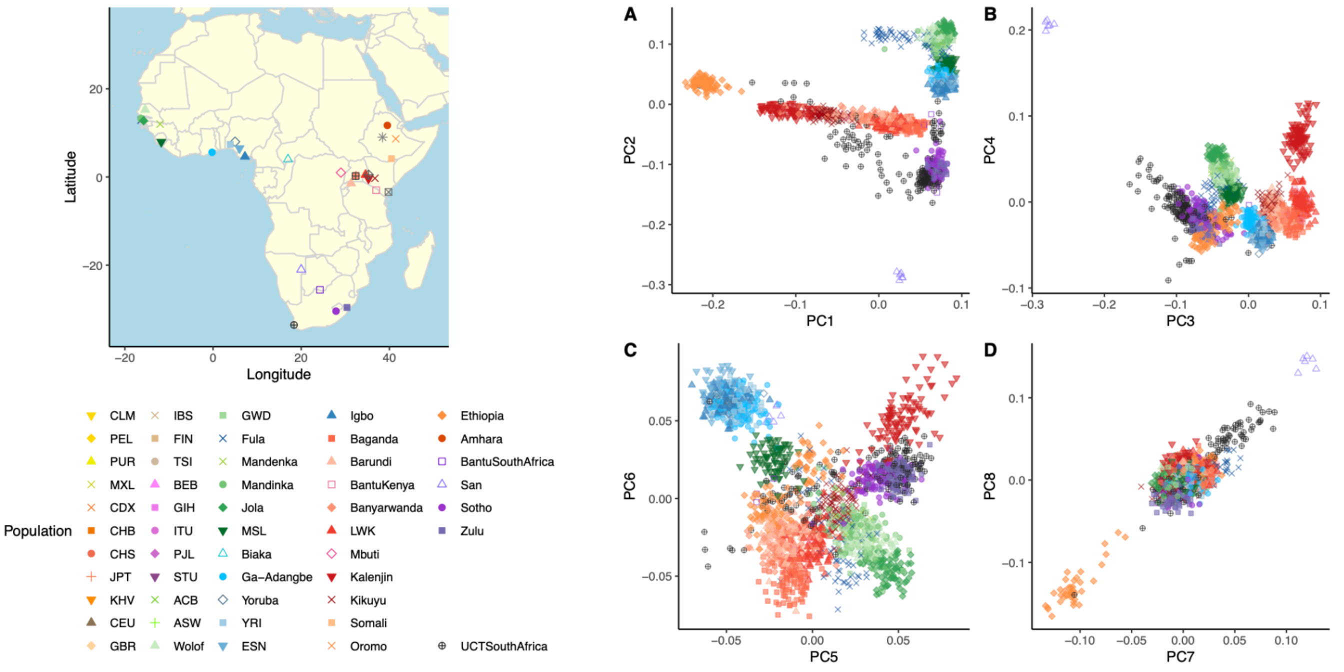


***Supplementary Figure 6***. *Fine-scale structure of genetic variation in the UCTSouthAfrica NeuroGAP-Psychosis collection site. A map showing the location of populations plotted is shown on the left.* ***A-D****) PCA plots for PCs 1-8 showing clustering of UCTSouthAfrica NeuroGAP-Psychosis samples across PC space with an African reference panel.*

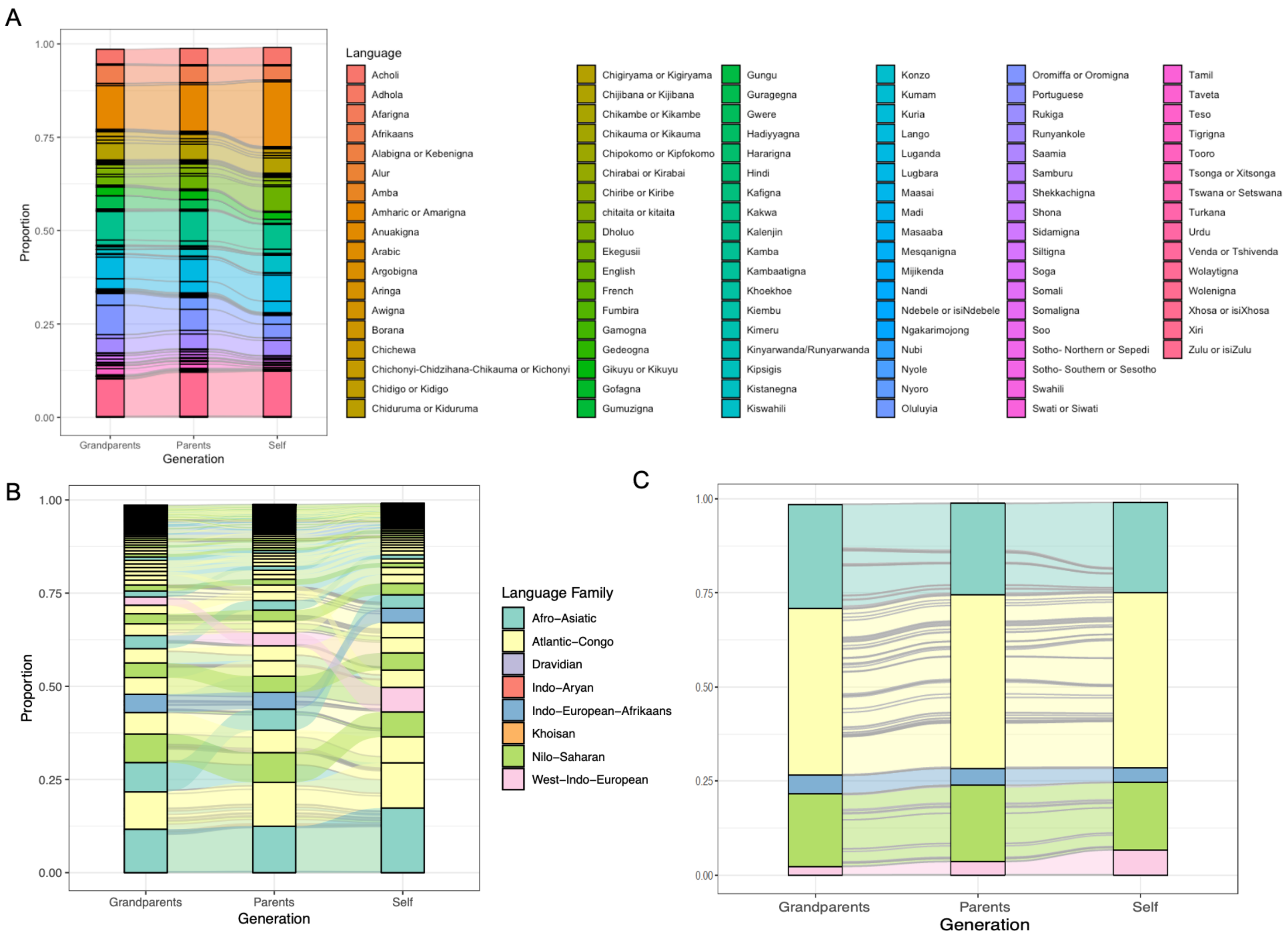


***Supplementary Figure 7.*** *Phenotypic composition of NeuroGAP-Psychosis samples. Alluvial plot showing the full self-reported primary language reports from participants. A) Primary languages shown individually across the pedigree. B) Primary languages sorted by frequency in each generation and colored by language family. C) Primary language frequency change over generations. Individual strata (separated by gray lines) show specific languages within each language group.*

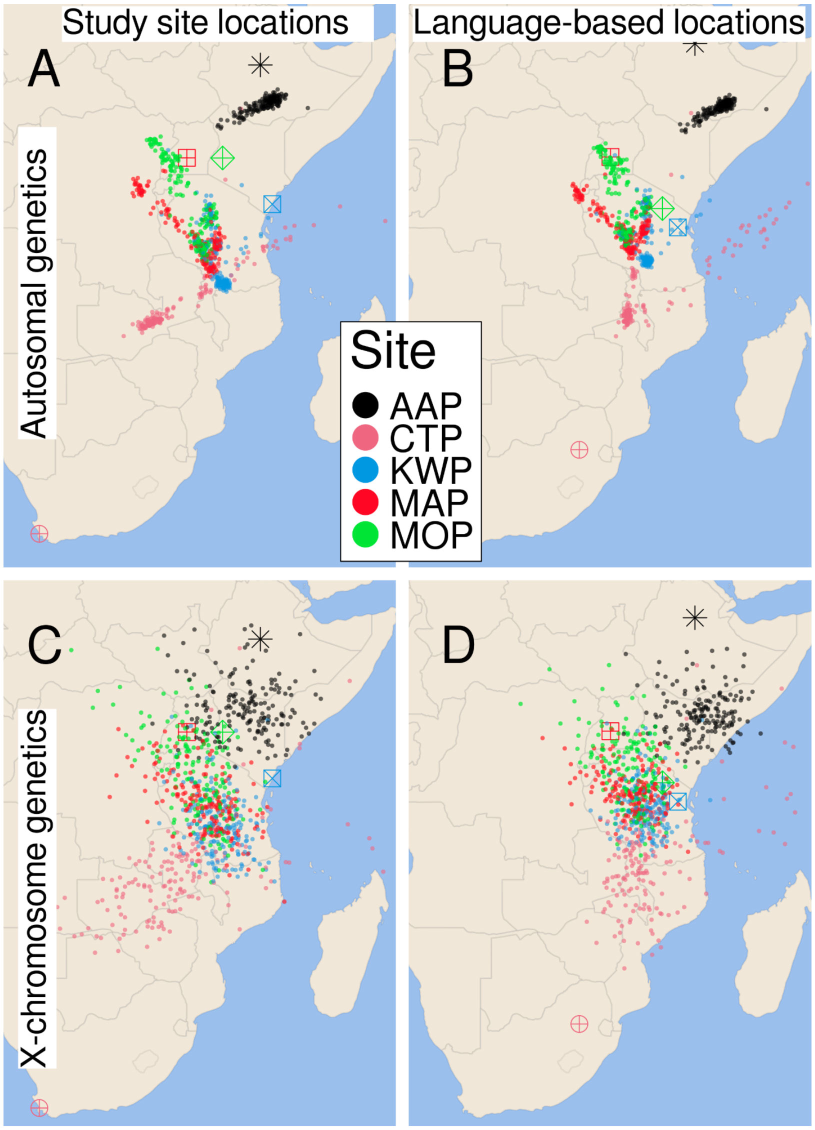


***Supplementary Figure 8.*** *Procrustes analyses indicate that autosomal genetic diversity is better correlated with geography than is X chromosome diversity. Plots represent the first three genetic PCs after a procrustes transformation. The upper panels use PCs generated using autosomal variation, and the lower panels use X chromosome variation. The left column uses the locations of the study site at which each individual was sampled; the right column uses each individual's self-reported languages and the centroids of these languages to identify a geographic midpoint of that individual's languages. Individuals are colored by primary field site. For each primary field site, the midpoint of individuals' locations (by study site or languages spoken) is represented by a large point.*

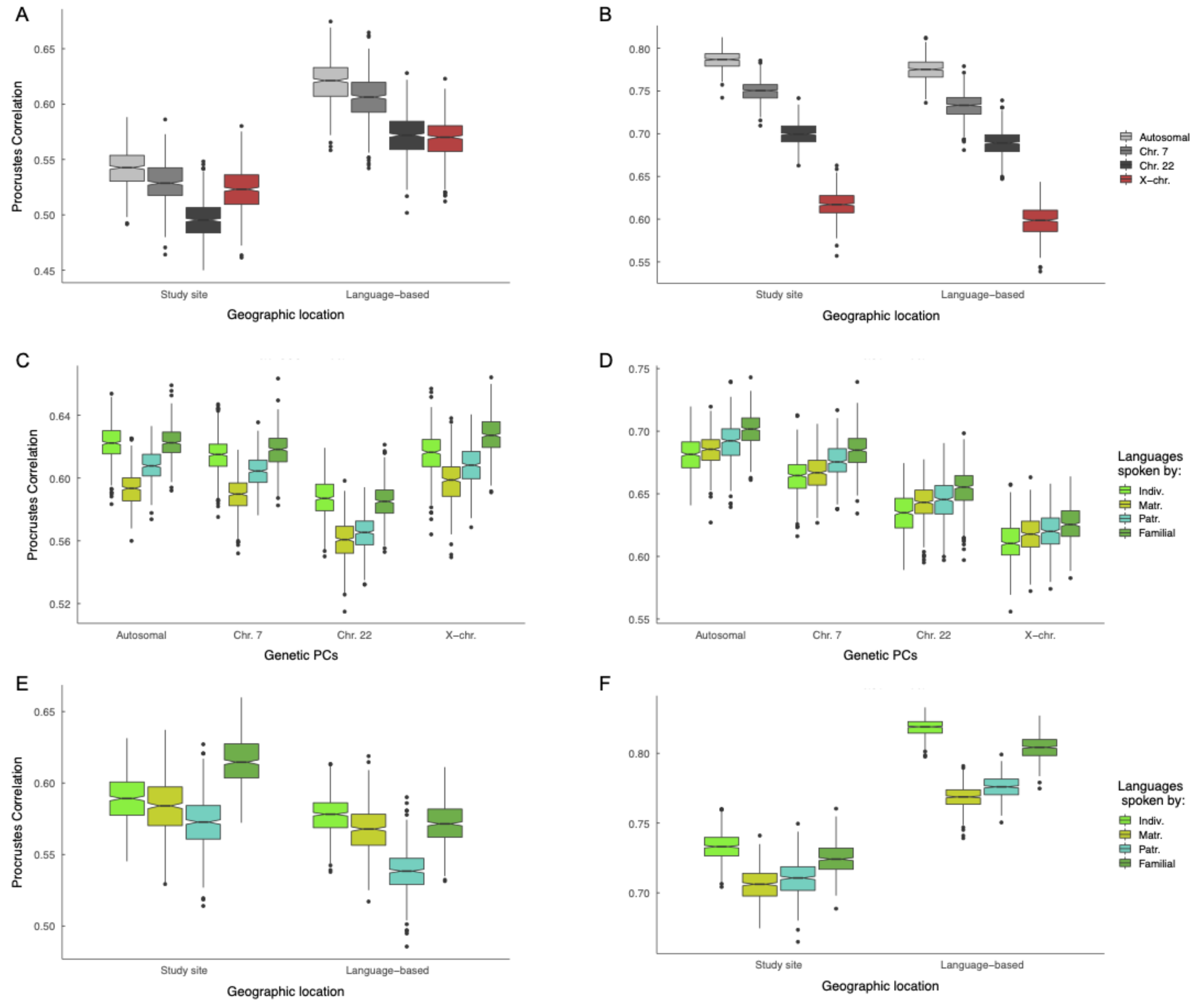


***Supplementary Figure 9***. *Procrustes correlations between genetics, geography, and language (all p < 5E-5). Procrustes correlations are shown between:* ***A,B****) geography and genetics.* ***C,D)*** *genetics and language, and* ***E,F)*** *geography and language. The left column includes results for the entire NeuroGAP collection. The right column contains results subset to the four cohorts in East Africa. Genetics analyses were conducted for the complete autosomes, the X chromosome, and two autosomal comparisons to the X: chr7 (similar length) and chr22 (similar SNP count). For linguistic analyses, linguistic variation is measured by the first three PCs of*

*phoneme inventories from languages reported by individuals as spoken by themselves and their relatives. Matrilineal relatives include the mother and maternal grandmother. Patrilineal relatives include the father and paternal grandfather. Familial refers to a weighted average of all reported family members. Note that Y-axis labels vary between plots.*

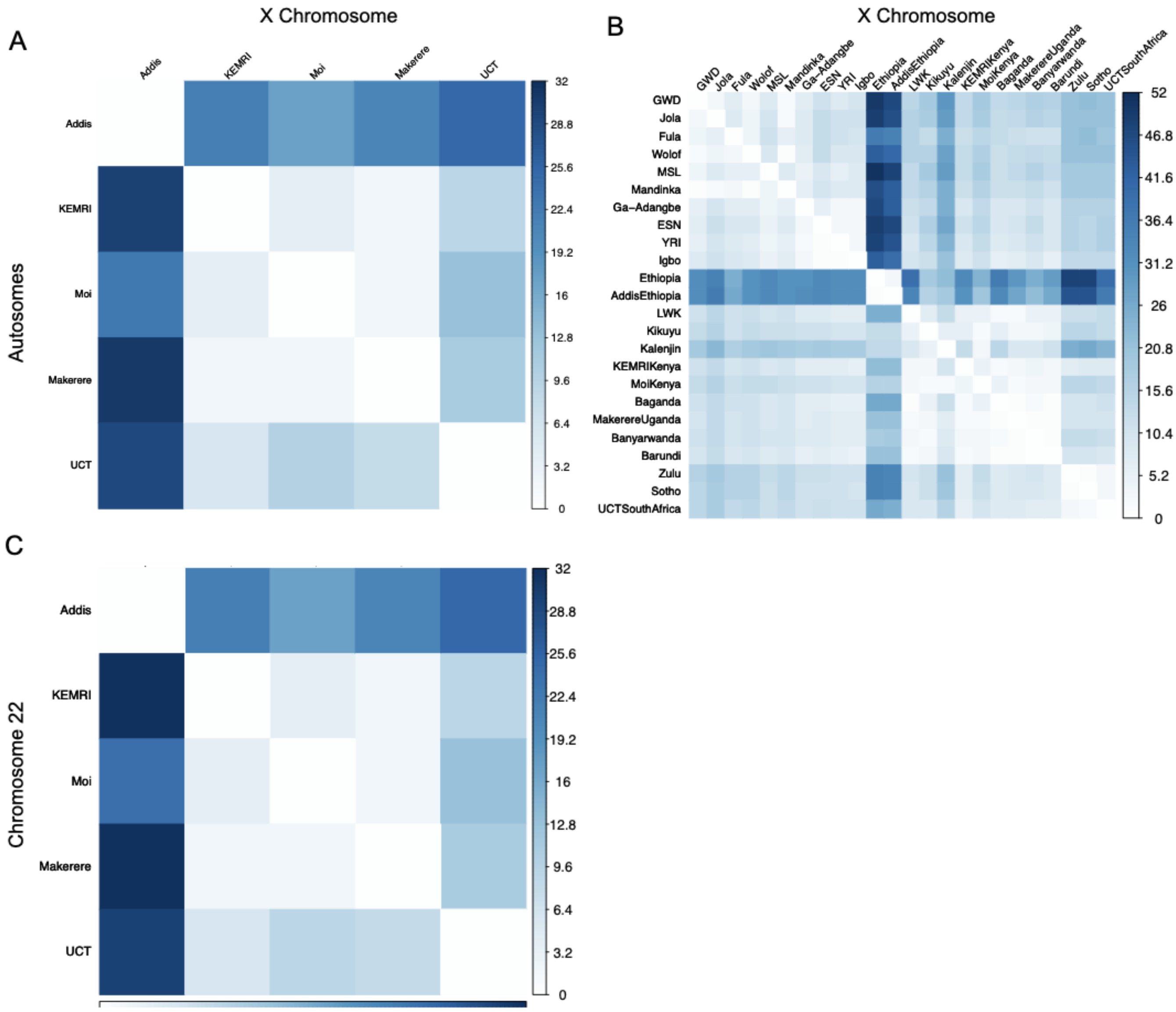

***Supplementary Figure 10.*** *Genetic differentiation across the autosomes compared to the X chromosome. Heatmap showing the $F_{ST}$ estimates calculated between pairwise populations' autosomes (below the diagonal) as compared to the X chromosome (above the diagonal). $F_{ST}$ values are multiplied by 1000 for easier interpretation.* ***A, C)*** *$F_{ST}$ estimates just between NeuroGAP-Psychosis collection sites. Panel A includes the entire autosomes while panel C is only chromosome 22 for comparison.* ***B)*** *$F_{ST}$ estimates between NeuroGAP-Psychosis collection sites as well as all African populations in our reference panel.*

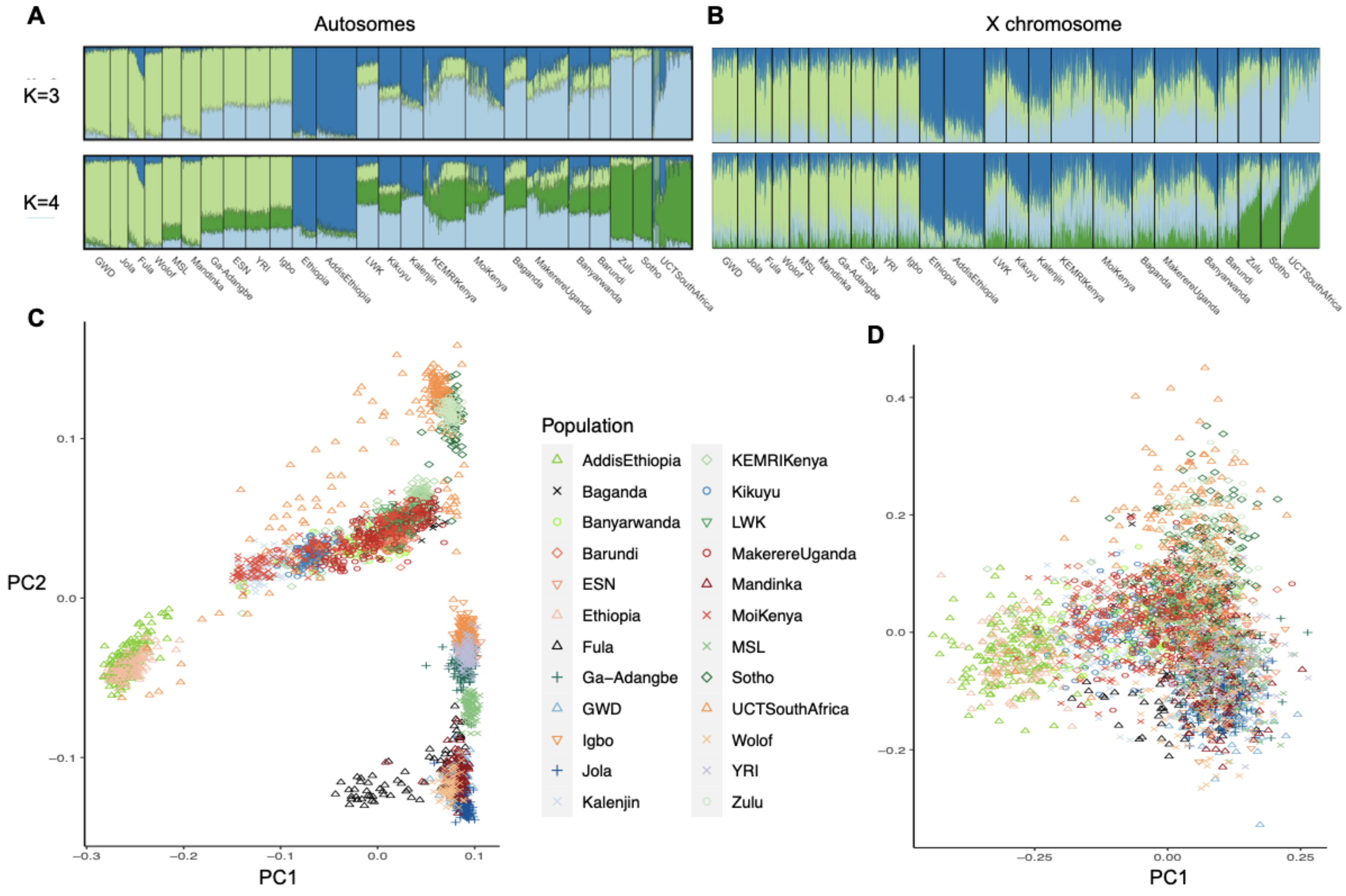

***Supplementary Figure 11***. *Comparison of ancestry proportions on the autosomes as compared to the X chromosome. Autosomes are shown in the left column, X chromosome on the right.* ***A-B****) ADMIXTURE runs at k=3 and 4. Colors are matched with light green tagging east African genetic variation, dark blue tagging Ethiopian variation, light blue tagging west African component, and dark green tagging a south African component.* ***C-D)*** *PC plots for the first two principal components of genetic variation in the autosomes and X chromosome.*

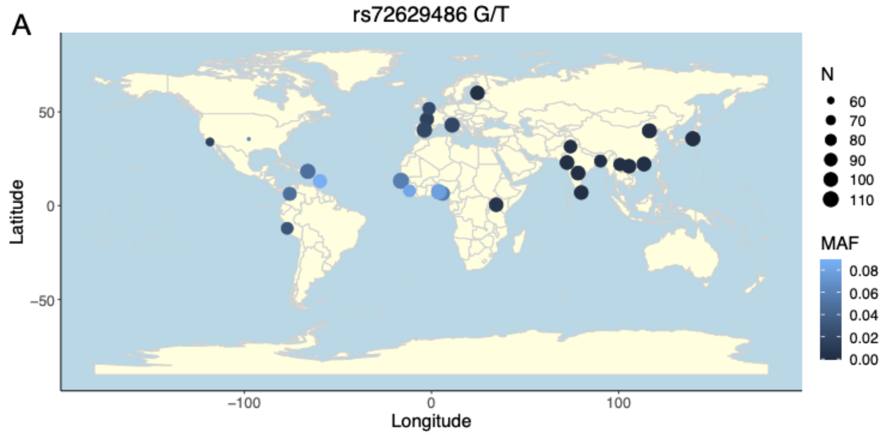


B

**Population Frequencies**

| Population | Allele Count | Allele Number | Number of Homozygotes | Allele Frequency |
|---|---|---|---|---|
| African/African-American | 1361 | 24624 | 40 | 0.05527 |
| Ashkenazi Jewish | 317 | 10294 | 4 | 0.03079 |
| Other | 170 | 7194 | 3 | 0.02363 |
| Latino/Admixed American | 766 | 35412 | 16 | 0.02163 |
| European (non-Finnish) | 1324 | 127584 | 11 | 0.01038 |
| South Asian | 139 | 30612 | 4 | 0.004541 |
| European (Finnish) | 58 | 25112 | 1 | 0.002310 |
| East Asian | 0 | 19890 | 0 | 0.000 |
| Female | 2018 | 127872 | 39 | 0.01578 |
| Male | 2117 | 152850 | 40 | 0.01385 |
| **Total** | **4135** | **280722** | **79** | **0.01473** |

***Supplementary Figure 12.*** *African genetic variation is broadly informative.* ***A)*** *the frequency of rs2071348, previously demonstrated to influence beta thalassemia, varies within the African continent dramatically, even across only our 5 pilot NeuroGAP-Psychosis sites. In Africa alone, missense variant rs72629486 spans the entire range of global frequencies reported in the gnomad database.* ***B)*** *Screenshot of the population frequencies of rs72629486 in gnomAD; Feb 28, 2021.*

**Supplementary Tables**

***Supplementary Table 1.*** Key for location/country and dataset of origin for all populations included in analyses. 'Region' indicates the continental assignment for non-African populations and the geographic region assigned within Africa as according to the UN Statistics division geoscheme (United Nations Statistics Division).

| Population | Cohort description/location | Dataset | Region | Language family (Africa) |
|---|---|---|---|---|
| CLM | Colombian in Medellín, Colombia | 1KG | AMR | |
| PEL | Peruvian in Lima Peru | 1KG | AMR | |
| PUR | Puerto Rican in Puerto Rico | 1KG | AMR | |
| MXL | Mexican Ancestry in Los Angeles CA USA | 1KG | AMR | |
| CDX | Chinese Dai in Xishuangbanna, China | 1KG | EAS | |
| CHB | Han Chinese in Beijing, China | 1KG | EAS | |
| CHS | Han Chinese South | 1KG | EAS | |
| JPT | Japanese in Tokyo, Japan | 1KG | EAS | |
| KHV | Kinh in Ho Chi Minh City, Vietnam | 1KG | EAS | |
| CEU | United States [EUR] | 1KG | EUR | |
| GBR | British From England and Scotland | 1KG | EUR | |
| IBS | Iberian Populations in Spain | 1KG | EUR | |
| FIN | Finnish in Finland | 1KG | EUR | |
| TSI | Toscani in Italia | 1KG | EUR | |
| BEB | Bengali in Bangladesh | 1KG | SAS | |
| GIH | Gujarati Indians in Houston, Texas, USA | 1KG | SAS | |
| ITU | Indian Telugu in the U.K. | 1KG | SAS | |
| PJL | Punjabi in Lahore, Pakistan | 1KG | SAS | |
| STU | Sri Lankan Tamil in the UK | 1KG | SAS | |
| ACB | African Caribbean in Barbados | 1KG | AFR | |
| ASW | African Ancestry in SW USA | 1KG | AFR | |

| | | | | |
|---|---|---|---|---|
| Wolof | Gambia | AGVP | West | Niger-Congo |
| GWD | Gambian in Western Division – Mandinka | 1KG | West | Niger-Congo |
| Mandenka | Mandenka in Senegal | HGDP | West | Niger-Congo |
| Mandinka | Gambia | AGVP | West | Niger-Congo |
| Jola | Gambia | AGVP | West | Niger-Congo |
| MSL | Mende in Sierra Leone | 1KG | West | Niger-Congo |
| Biaka | Biaka in Central African Republic | HGDP | Middle | Niger-Congo |
| Ga-Adangbe | Ghana | AGVP | West | Niger-Congo |
| Yoruba | Yoruba in Nigeria | AGVP | West | Niger-Congo |
| YRI | Yoruba in Ibadan, Nigeria | 1KG | West | Niger-Congo |
| ESN | Esan in Nigeria | 1KG | West | Niger-Congo |
| Igbo | Nigeria | AGVP | West | Niger-Congo |
| Fula | Gambia | AGVP | West | Niger-Congo |
| Baganda | Uganda | AGVP | East | Niger-Congo |
| Barundi | Uganda | AGVP | East | Niger-Congo |
| BantuKenya | Bantu in Kenya | HGDP | East | Niger-Congo |
| Banyarwanda | Uganda | AGVP | East | Niger-Congo |
| LWK | Luhya in Webuye, Kenya | 1KG | East | Niger-Congo |
| Mbuti | Mbuti in Democratic Republic of Congo | HGDP | Middle | Niger-Congo |
| Kalenjin | Kenya | AGVP | East | Nilo-Saharan |
| Kikuyu | Kenya | AGVP | East | Niger-Congo |
| Somali | Ethiopia | AGVP | East | Afro-Asiatic |
| Oromo | Ethiopia | AGVP | East | Afro-Asiatic |
| Ethiopia | Ethiopia | AGVP | East | Afro-Asiatic |
| Amhara | Ethiopia | AGVP | East | Afro-Asiatic |
| BantuSouthAfrica | Bantu in South Africa | HGDP | South | Niger-Congo |
| San | San in Namibia | HGDP | South | Khoisan |
| Sotho | South Africa | AGVP | South | Niger-Congo |
| Zulu | South Africa | AGVP | South | Niger-Congo |
| AddisEthiopia | Ethiopia | NeuroGAP | East | Assortment |
| KEMRIKenya | Kenya | NeuroGAP | East | Assortment |

| | | | | | |
|---|---|---|---|---|---|
| MoiKenya | Kenya | | NeuroGAP | East | Assortment |
| MakerereUganda | Uganda | | NeuroGAP | East | Assortment |
| UCTSouthAfrica | South Africa | | NeuroGAP | South | Assortment |

***Supplementary Table 2.*** *Raw data for language phenotypes reported for each familial relationship across the NeuroGAP-Psychosis dataset. See STAR Methods section 'Ethnolinguistic Phenotypes' for a detailed description of the specific phenotypes collected.*

| Language | lang_self_1 | lang_mat_1 | lang_pat_1 | lang_mgm_1 | lang_pgf_1 | lang_pgm_1 | lang_mgf_1 | Language |
|---|---|---|---|---|---|---|---|---|
| 1 | 819 | 764 | 737 | 576 | 379 | 495 | 413 | Acholi |
| 2 | 49 | 49 | 46 | 47 | 35 | 38 | 42 | Adhola |
| 4 | 680 | 820 | 746 | 706 | 460 | 579 | 571 | Afrikaans |
| 5 | 1 | 2 | 2 | 0 | 0 | 0 | 0 | Alabigna or Kebenigna |
| 6 | 84 | 100 | 95 | 82 | 57 | 67 | 73 | Alur |
| 8 | 3085 | 2210 | 2098 | 1545 | 1230 | 1334 | 1426 | Amharic or Amarigna |
| 9 | 1 | 0 | 0 | 0 | 0 | 0 | 1 | Anuakigna |
| 10 | 37 | 39 | 72 | 26 | 40 | 38 | 27 | Arabic |
| 12 | 31 | 46 | 52 | 34 | 21 | 31 | 25 | Aringa |
| 13 | 3 | 5 | 5 | 9 | 7 | 6 | 7 | Awigna |
| 16 | 8 | 7 | 8 | 5 | 3 | 6 | 3 | Borana |
| 18 | 189 | 195 | 197 | 165 | 120 | 152 | 144 | Chichonyi-Chidzihana-Chikauma or Kichonyi |
| 19 | 126 | 132 | 143 | 134 | 105 | 122 | 102 | Chidigo or Kidigo |
| 20 | 116 | 120 | 121 | 92 | 97 | 110 | 99 | Chiduruma or Kiduruma |
| 21 | 725 | 717 | 716 | 631 | 446 | 564 | 486 | Chigiryama or Kigiryama |
| 22 | 38 | 25 | 38 | 21 | 24 | 27 | 19 | Chijibana or Kijibana |

| Language | lang_self_1 | lang_mat_1 | lang_pat_1 | lang_mgm_1 | lang_pgf_1 | lang_pgm_1 | lang_mgf_1 | Language |
|---|---|---|---|---|---|---|---|---|
| 23 | 22 | 20 | 24 | 25 | 15 | 17 | 15 | Chikambe or Kikambe |
| 24 | 52 | 55 | 57 | 44 | 34 | 47 | 39 | Chikauma or Kikauma |
| 25 | 27 | 34 | 28 | 28 | 11 | 20 | 22 | Chipokomo or Kipfokomo |
| 26 | 51 | 54 | 59 | 41 | 37 | 47 | 39 | Chirabai or Kirabai |
| 27 | 5 | 6 | 8 | 5 | 4 | 4 | 2 | Chiribe or Kiribe |
| 28 | 155 | 193 | 146 | 158 | 92 | 105 | 129 | chitaita or kitaita |
| 29 | 209 | 260 | 273 | 219 | 164 | 214 | 163 | Dholuo |
| 30 | 78 | 116 | 117 | 92 | 69 | 94 | 68 | Ekegusii |
| 31 | 1173 | 569 | 619 | 299 | 224 | 265 | 256 | English |
| 33 | 16 | 13 | 21 | 10 | 5 | 7 | 7 | French |
| 34 | 20 | 29 | 38 | 27 | 20 | 25 | 27 | Fumbira |
| 36 | 12 | 32 | 30 | 34 | 33 | 34 | 34 | Gamogna |
| 37 | 1 | 3 | 2 | 1 | 1 | 1 | 1 | Gedeogna |
| 38 | 340 | 426 | 381 | 354 | 197 | 266 | 271 | Gikuyu or Kikuyu |
| 39 | 1 | 3 | 4 | 4 | 2 | 3 | 1 | Gofagna |
| 41 | 1 | 0 | 1 | 0 | 1 | 0 | 0 | Gungu |
| 42 | 19 | 24 | 22 | 29 | 20 | 29 | 19 | Gwere |
| 43 | 17 | 31 | 33 | 38 | 33 | 35 | 32 | Hadiyyagna |
| 46 | 4 | 4 | 5 | 4 | 5 | 4 | 3 | Kafigna |
| 47 | 24 | 33 | 32 | 25 | 17 | 25 | 16 | Kakwa |
| 48 | 1188 | 1391 | 1369 | 1073 | 775 | 963 | 822 | Kalenjin |
| 49 | 198 | 237 | 203 | 193 | 133 | 159 | 151 | Kamba |
| 50 | 10 | 17 | 12 | 13 | 12 | 13 | 13 | Kambaatigna |
| 52 | 11 | 13 | 11 | 12 | 9 | 8 | 10 | Kiembu |
| 53 | 43 | 50 | 44 | 48 | 27 | 33 | 38 | Kimeru |
| 54 | 8 | 14 | 17 | 9 | 7 | 7 | 4 | Kipsigis |
| 56 | 825 | 317 | 317 | 151 | 109 | 142 | 122 | Kiswahili |
| 57 | 17 | 19 | 22 | 18 | 16 | 18 | 14 | Konzo |

| Language | lang_self_1 | lang_mat_1 | lang_pat_1 | lang_mgm_1 | lang_pgf_1 | lang_pgm_1 | lang_mgf_1 | Language |
|---|---|---|---|---|---|---|---|---|
| 59 | 14 | 17 | 11 | 14 | 11 | 10 | 13 | Kumam |
| 61 | 7 | 10 | 9 | 6 | 5 | 7 | 6 | Kupsapiiny |
| 62 | 2 | 4 | 4 | 4 | 2 | 2 | 3 | Kuria |
| 63 | 87 | 102 | 106 | 92 | 76 | 82 | 74 | Lango |
| 64 | 1242 | 1066 | 1012 | 765 | 604 | 703 | 672 | Luganda |
| 65 | 544 | 536 | 508 | 381 | 266 | 328 | 307 | Lugbara |
| 66 | 10 | 25 | 19 | 23 | 11 | 18 | 19 | Ma'di |
| 67 | 13 | 19 | 17 | 13 | 16 | 20 | 10 | Maasai |
| 68 | 32 | 49 | 48 | 49 | 38 | 49 | 40 | Masaaba |
| 69 | 1 | 0 | 0 | 0 | 0 | 0 | 0 | Mesqanigna |
| 70 | 1 | 1 | 1 | 1 | 1 | 1 | 1 | Mijikenda |
| 71 | 28 | 42 | 41 | 24 | 23 | 25 | 21 | Nandi |
| 72 | 15 | 17 | 10 | 18 | 7 | 8 | 6 | Ndebele or isiNdebele |
| 74 | 2 | 5 | 2 | 4 | 3 | 4 | 2 | Ng'akarimojong |
| 76 | 6 | 8 | 15 | 8 | 8 | 8 | 5 | Nubi |
| 78 | 11 | 14 | 20 | 12 | 15 | 17 | 14 | Nyole |
| 79 | 28 | 43 | 53 | 33 | 28 | 35 | 29 | Nyoro |
| 80 | 422 | 556 | 529 | 438 | 324 | 390 | 347 | Oluluyia |
| 82 | 647 | 950 | 1000 | 980 | 898 | 949 | 907 | Oromiffa or Oromigna |
| 84 | 5 | 8 | 6 | 5 | 3 | 4 | 5 | Pokot |
| 85 | 3 | 5 | 6 | 4 | 3 | 3 | 2 | Portuguese |
| 87 | 723 | 703 | 676 | 510 | 413 | 466 | 448 | Runyankole |
| 88 | 22 | 23 | 28 | 21 | 20 | 19 | 18 | Saamia |
| 89 | 1 | 1 | 2 | 0 | 2 | 1 | 0 | Samburu |
| 91 | 1 | 1 | 1 | 1 | 2 | 2 | 1 | Shekkachigna |
| 93 | 12 | 16 | 18 | 19 | 18 | 16 | 19 | Sidamigna |
| 94 | 66 | 124 | 127 | 110 | 110 | 104 | 107 | Siltigna |
| 95 | 107 | 129 | 129 | 100 | 96 | 114 | 88 | Soga |
| 96 | 35 | 42 | 38 | 26 | 22 | 25 | 16 | Somali |

| Language | lang_self_1 | lang_mat_1 | lang_pat_1 | lang_mgm_1 | lang_pgf_1 | lang_pgm_1 | lang_mgf_1 | Language |
|---|---|---|---|---|---|---|---|---|
| 99 | 7 | 4 | 13 | 9 | 5 | 7 | 6 | Sotho- Northern or Sepedi |
| 100 | 67 | 79 | 91 | 63 | 30 | 52 | 54 | Sotho- Southern or Sesotho |
| 101 | 31 | 39 | 49 | 24 | 20 | 22 | 17 | Swahili |
| 102 | 3 | 5 | 5 | 2 | 3 | 3 | 5 | Swati or Siwati |
| 104 | 4 | 4 | 4 | 5 | 2 | 2 | 3 | Taveta |
| 105 | 102 | 122 | 126 | 111 | 85 | 97 | 109 | Teso |
| 106 | 69 | 190 | 201 | 203 | 179 | 191 | 193 | Tigrigna |
| 107 | 31 | 53 | 31 | 45 | 18 | 30 | 36 | Tooro |
| 108 | 16 | 18 | 18 | 10 | 5 | 9 | 5 | Tsonga or Xitsonga |
| 110 | 9 | 13 | 18 | 10 | 4 | 8 | 3 | Tswana or Setswana |
| 111 | 13 | 18 | 19 | 13 | 8 | 8 | 11 | Turkana |
| 113 | 4 | 6 | 7 | 2 | 1 | 1 | 4 | Venda or Tshivenda |
| 114 | 21 | 41 | 35 | 47 | 47 | 43 | 49 | Wolaytigna |
| 115 | 25 | 47 | 41 | 37 | 26 | 31 | 31 | Wolenigna |
| 116 | 2154 | 2105 | 1970 | 1589 | 858 | 1204 | 1114 | Xhosa or isiXhosa |
| 117 | 2 | 1 | 0 | 0 | 0 | 0 | 0 | Xiri |
| 119 | 44 | 43 | 56 | 37 | 21 | 36 | 18 | Zulu or isiZulu |
| 120 | 191 | 439 | 450 | 453 | 387 | 426 | 389 | Guragegna |
| 121 | 121 | 163 | 163 | 129 | 98 | 118 | 107 | Rukiga |
| 122 | 26 | 78 | 60 | 97 | 46 | 72 | 78 | Kinyarwanda/Runyarwanda |
| 123 | 93 | 94 | 99 | 58 | 44 | 64 | 50 | Shona |
| 124 | 32 | 30 | 28 | 19 | 7 | 11 | 10 | Chichewa |
| 999 | 113 | 186 | 180 | 167 | 117 | 139 | 129 | Other (Describe) |
| -777 | 0 | 4 | 0 | 5 | 3 | 2 | 2 | Unknown |
| -666 | 0 | 2 | 1 | 1 | 1 | 0 | 1 | NA |
| 7 | 0 | 1 | 1 | 1 | 1 | 0 | 0 | Amba |
| 40 | 0 | 1 | 1 | 0 | 0 | 0 | 0 | Gumuzigna |

| Language | lang_self_1 | lang_mat_1 | lang_pat_1 | lang_mgm_1 | lang_pgf_1 | lang_pgm_1 | lang_mgf_1 | Language |
|---|---|---|---|---|---|---|---|---|
| 44 | 0 | 4 | 5 | 2 | 2 | 1 | 1 | Hararigna |
| 55 | 0 | 1 | 1 | 1 | 2 | 1 | 1 | Kistanegna |
| 98 | 0 | 2 | 0 | 0 | 0 | 0 | 0 | Soo |
| 103 | 0 | 1 | 0 | 0 | 0 | 0 | 0 | Tamil |
| 3 | 0 | 0 | 1 | 0 | 0 | 0 | 0 | Afarigna |
| 11 | 0 | 0 | 1 | 0 | 0 | 0 | 0 | Argobigna |
| 45 | 0 | 0 | 1 | 0 | 0 | 0 | 0 | Hindi |
| 51 | 0 | 0 | 0 | 1 | 1 | 1 | 1 | Khoekhoe |
| 97 | 0 | 0 | 0 | 4 | 2 | 1 | 0 | Somaligna |
| 112 | 0 | 0 | 0 | 1 | 1 | 0 | 1 | Urdu |

***Supplementary Table 3**. Classification of self-reported primary ethnicities included in the surveys, with associated data collected from the Ethnographic Atlas. See STAR Methods section 'Ethnolinguistic Phenotypes' for a detailed description of the specific phenotypes collected.*

| Ethnicity | Ethnographic Atlas ID | Inheritance pattern according to the Ethnographic Atlas (EA076) |
|---|---|---|
| Afar | Ca6 | NA |
| Alaba-K'abeena | NA | NA |
| Amhara | Ca7 | Patrilineal |
| Anuak | Ai44 | NA |
| Argobba | NA | NA |
| Aringa | NA | NA |
| Awngi | NA | NA |
| Babwisi | Ad13 | NA |
| Bafumbira | NA | NA |

| Ethnicity | Ethnographic Atlas ID | Inheritance pattern according to the Ethnographic Atlas (EA076) |
|---|---|---|
| Baganda | Ad7 | Patrilineal |
| Bagisu | Ad9 | Patrilineal |
| Bagwere | NA | NA |
| Bahororo | NA | NA |
| Bakiga | Ad13 | Patrilineal |
| Bakonzo | Ad44 | NA |
| Banyankore | Ad45 | Patrilineal |
| Banyarwanda | Ae10 | Patrilineal |
| Banyole | NA | NA |
| Banyoro | Ad2 | Patrilineal |
| Bapedi North Sotho | Ab15 | Patrilineal |
| Baruli | NA | NA |
| Basamia | NA | NA |
| Basoga | Ad46 | Patrilineal |
| Basotho South Sotho | Ab8 | Patrilineal |
| Basuba | NA | NA |
| Batoro | Ad48 | NA |
| Borana | Ca11 | Patrilineal |
| Coloured Cape Malay | Ej8 | Neither |
| Coloured Khoisan | NA | NA |
| Coloured Other | NA | NA |
| Dodoth | Aj30 | NA |
| Embu | NA | NA |
| Gabra | NA | NA |
| Gamo | NA | NA |
| Gedeo | Ca15 | Patrilineal |
| Gofa | NA | NA |

| Ethnicity | Ethnographic Atlas ID | Inheritance pattern according to the Ethnographic Atlas (EA076) |
|---|---|---|
| Gumuz | NA | NA |
| Hadiyya | NA | NA |
| Harari | NA | NA |
| Indian or Asian | NA | NA |
| Iteso | Aj1 | Patrilineal |
| Jie | Aj21 | Patrilineal |
| Jonam | Aj17 | Patrilineal |
| Jopadhola | Aj10 | Patrilineal |
| Kafa | Ka30 | NA |
| Kakwa | Aj14 | NA |
| Kalenjin | Aj9 | Patrilineal |
| Kamba | Ad34 | Patrilineal |
| Kambaata | NA | NA |
| Karimojong | Aj30 | NA |
| Kenyan Somali | Ca2 | Neither |
| Kikuyu | Ad4 | Patrilineal |
| Kisii | Ad12 | Patrilineal |
| Kistane | Ca8 | Patrilineal |
| Kumam | NA | NA |
| Kuria | NA | NA |
| Langi | Ad25 | Matrilineal |
| Lugbara | Ai32 | NA |
| Luhya | Ad41 | Patrilineal |
| Luo | Aj6 | Patrilineal |
| Maasai | Aj2 | Patrilineal |
| Madi | Ai33 | Patrilineal |
| Mbeere | NA | NA |

| Ethnicity | Ethnographic Atlas ID | Inheritance pattern according to the Ethnographic Atlas (EA076) |
|---|---|---|
| Meru | Ad35 | Patrilineal |
| Mesqan | Ca8 | Patrilineal |
| Mijikenda | Ad31 | Matrilineal |
| Ndebele | Ab9 | Patrilineal |
| Nuer | Aj3 | Patrilineal |
| Orma | Ca12 | Patrilineal |
| Oromo | Ca12 | Patrilineal |
| Other African | NA | NA |
| Pokot | Aj23 | Patrilineal |
| Rendile | NA | NA |
| Sabiny | Aj27 | NA |
| Samburu | Aj29 | Patrilineal |
| Sebat Bet Gurage | Ca8 | Patrilineal |
| Shekkacho | NA | NA |
| Sheko | Ka42 | NA |
| Sidama | Ca16 | Patrilineal |
| Silt'e | NA | NA |
| Somali | Ca2 | Neither |
| Swahili | NA | NA |
| Swazi | Ab2 | Patrilineal |
| Taita | Ad37 | Patrilineal |
| Teso | Aj1 | Patrilineal |
| Tharaka | Ad35 | Patrilineal |
| Tigrigna | Ca3 | Patrilineal |
| Tsonga | Ab4 | Patrilineal |
| Tswana | Ab13 | Patrilineal |
| Turkana | Aj5 | Patrilineal |

| Ethnicity | Ethnographic Atlas ID | Inheritance pattern according to the Ethnographic Atlas (EA076) |
|---|---|---|
| Venda | Ab6 | Patrilineal |
| White - Afrikaner | NA | NA |
| White - Other | NA | NA |
| Wolane | Ca8 | Patrilineal |
| Wolaytta | NA | NA |
| Xhosa | Ab11 | Patrilineal |
| Zulu | Ab12 | Patrilineal |
| Other (please specify) | NA | NA |
| Unknown | NA | NA |
| Sodo Gurage | Ca8 | Patrilineal |
| Acholi-Luo | Aj6 | Patrilineal |
| Alur | Aj17 | Patrilineal |
| Giriama | Ad32 | Patrilineal |
| Digo | Ad30 | Matrilineal |
| Chonyi | Ad31 | Matrilineal |
| Duruma | Ad31 | Matrilineal |
| Jibana | Ad31 | Matrilineal |
| Kambe | Ad34 | Patrilineal |
| Kauma | Ad31 | Matrilineal |
| Rabai | Ad31 | Matrilineal |
| Ribe | Ad31 | Matrilineal |
| Shona | Ab18 | Patrilineal |
| Pokomo | Ad33 | Patrilineal |
| Arab | Cb13 | NA |

***Supplementary Table 4***. *Variant and individual counts throughout the Autosomal QC process.*

| **Autosomal QC Filter Results** | | | | | |
|---|---|---|---|---|---|
| **Filter Name** | **Variants or Individuals Remaining After Filter per Pilot Site** | | | | |
| | Moi, Kenya | Ethiopia | KEMRI | South Africa | Uganda |
| Autocall Call Rate (samples) | 189 | 183 | 188 | 185 | 192 |
| Variant Call Rate (variants) | 638235 | 638235 | 638235 | 638235 | 638235 |
| Individual Call Rate (samples) | 189 | 181 | 188 | 182 | 190 |
| Sex Violations (samples) | 187 | 181 | 188 | 179 | 188 |
| Minor Allele Frequency (variants) | 360321 | 360321 | 360321 | 360321 | 360321 |
| Hardy Weinberg Equilibrium (variants) | 331667 | 331667 | 331667 | 331667 | 331667 |
| Sample Relatedness (samples) | 173 | 179 | 187 | 175 | 186 |
| **Final Counts (variants / samples)** | 331667 / 173 | 331667 / 179 | 331667 / 187 | 331667 / 175 | 331667 / 186 |

***Supplementary Table 5***. *Variant counts throughout X Chromosome QC.*

| X Chromosome Variant QC Filter Results | | |
|---|---|---|
| **Filter Name** | **Variants Remaining After Filter** | |
| | PAR Region | Female nonPAR Region |
| Variant Call Rate | 515 | 16261 |
| MAF | 411 | 11113 |
| HWE | 402 | 11104 |
| **Final Counts** | 900 Samples<br>402 Variants | 900 Samples<br>11104 Variants |
